# Promoter architecture links gene duplication with transcriptional divergence

**DOI:** 10.1101/2021.07.03.450995

**Authors:** Evgeny Fraimovitch, Tzachi Hagai

## Abstract

Gene duplication is thought to be a central mechanism in evolution to gain new functions, but gene families vary greatly in their rates of gene duplication and long-term retention. Here, we discover a link between the promoter architecture of vertebrate genes and their rate of duplication: Genes that harbor CpG Islands in their promoters (CGI genes) – nearly 60% of our genes – have rarely duplicated in recent evolutionary times, and most CGI gene duplication events predate the emergence of CGI as a major regulatory element of vertebrate genes. In contrast, CGI-less genes predominate duplications that have occurred since the divergence of vertebrates. Furthermore, CGI-less paralogs are transcriptionally more divergent than CGI paralogs, even when comparing CGI and CGI-less paralogs that have duplicated at similar evolutionary times – suggesting greater capacity of CGI-less promoters to enable divergence in expression. This higher divergence between CGI-less paralogs is also reflected in lower similarity of transcription factors that bind to the promoters of CGI-less paralog pairs in comparison with CGI paralogs. Finally, CGI-less paralogs have a greater tendency to sub- and neo-functionalize, and they transcriptionally diversify faster following duplication. Our results highlight the links between promoter architecture, gene expression plasticity and their impact on gene expansion, and unravel an unappreciated role of CGI elements in shaping genome evolution.

**Significance statement:** Gene duplication generates extra gene copies, providing material for evolution of new functions. However, many duplicated genes are eliminated due to functional and regulatory constraints. The evolutionary processes that govern the elimination and persistence of duplicated genes are not well understood. Here, we focus on CpG Islands (CGIs) – important elements that occur in the majority of gene promoters. We show that genes with CGIs in their promoters have duplicated almost exclusively in ancient times, and nearly all recent duplications involve CGI-less genes. Furthermore, duplicated CGI-less genes diverge more in expression and display more distinctive transcription and cis-regulation compared to duplicated CGI-genes. Our results demonstrate how promoter structure influences transcriptional evolvability and, in turn, the retention of new genes.

## Introduction

Gene duplication introduces new gene copies and, as such, plays a central role in genome evolution and organismal complexity(1–5). Many of the duplicated genes, however, are not fixed following duplication and do not evolve into a functional paralog(5, 6). Successful gene retention is influenced by various factors, including dosage balance constraints(7), the mode of gene duplication – whole genome versus small-scale duplication(8), as well as gene evolvability and inter-paralog interactions(9, 10). As a result, retained genes are enriched for specific functions and genetic properties(11), but the causal mechanisms that lead to specific gene retention are not yet fully understood.

Gene duplication and fixation rates have been associated with gene expression levels as well as with the number of transcription initiation events(12–14), both of which are influenced by the structure of the gene’s promoter(15). However, the relationship between types of promoters and gene duplication has not been directly studied and how promoter architecture shapes retention and evolvability of genes following duplication remains largely unknown.

In mammals, more than half of the promoters of coding genes are associated with regions of non-methylated DNA, called CpG islands (**CGIs**), where CpG dinucleotide frequency is higher in comparison with other regions along the genome(16). These CGIs are thought to enable a transcriptionally permissive chromatin environment(15, 17) that opposes the repressive effects of methylation(18). CGI-rich promoters constitute a major class of promoters that have a characteristic chromatin organization and that is linked with specific patterns of transcription initiation and gene expression(19, 20). Genes with promoters containing CGIs (denoted as **CGI genes**) encode, among others, housekeeping genes expressed in numerous tissues(21, 22), developmental master regulators(23, 24), as well as genes that are expressed specifically in the central nervous system(25).

Here, we study the association between promoter architecture, the rate of gene duplication during vertebrate evolution, and the transcriptional divergence between the resulting duplicated genes. We focus on CGI as a major regulatory element in vertebrate promoters and analyze how patterns of gene duplication and transcriptional divergence between paralogs are linked to the presence of CGI regions in gene promoters. For this, we contrast CGI genes with genes lacking these CGI regions in their promoters (denoted as **CGI-less genes**), and show that the two classes differ significantly: (1) in the rate of gene duplication and retention in the course of vertebrate evolution, (2) in the level of transcriptional divergence between the resulting duplicates and in their ability to form sub/neo-functionalized copies, as well as (3) in the differences in transcription factors that bind to the promoters of these duplicated genes. Our study thus highlights an unappreciated aspect of gene evolvability, transcriptional divergence and neofunctionalization, and how they are shaped by promoter structures.

## Results

### Defining CGI and CGI-less genes

In this work we focus on protein-coding genes and use gene homology relationships and genome locations as annotated in ENSEMBL version 98(26). Since there is no well accepted method to define genes as harboring CGI in their promoters (**CGI genes**), we used two different approaches to define CGI genes: (1) A conservative approach, where genes are categorized as CGI genes if at least 50% of the region 300bp upstream of the transcription start site (**TSS**) and 100bp downstream of it overlaps with an annotated CGI region (following previous work by us and others(27, 28)). We treat all other genes as **CGI-less genes** – genes without a significant overlap of CGIs with their promoter region. (2) A more inclusive approach where any gene with at least 1bp overlap with a CGI in the region 1,000bp upstream and 1,000bp downstream of its TSS is categorized as a CGI gene (following other work(29)). We did not observe significant differences between the two approaches and report our results using the conservative approach in the main figures, and include, for comparison, some of the results based on the inclusive approach in Supporting Information. In humans, there are 11,230 CGI genes and 8,506 CGI-less genes (∼56.9% and ∼43.1%, respectively), while in mouse there are 12,527 CGI and 9,347 and CGI-less genes (57.2% and 42.8%, respectively), according to the conservative approach.

### CGI genes display low duplication rates across vertebrates

To study the rate of gene duplication in the two classes of genes – CGI and CGI-less genes – we first investigated the conservation of ortholog occurrence of these two gene classes across species. The occurrence of one-to-one orthologs across species can indicate low rate of gene duplication and loss. We calculated the fraction of human genes that have pairwise one-to-one orthologs with a group of 23 representative vertebrates and observed that CGI genes have a higher fraction of these orthologs in comparison with CGI-less genes in all tested species (FDR-corrected P-values in all comparisons of paired species were below 10^−5^, **Figs 1A** and **Supp Fig 2**). In agreement with this, human CGI-genes have a significantly higher portion of one-to-one orthologs among all annotated species in ENSEMBL, in comparison with CGI-less genes (P-value=1.6×10^−23^, one-sided Mann-Whitney test, **Fig 1B**).

**Figure 1:**
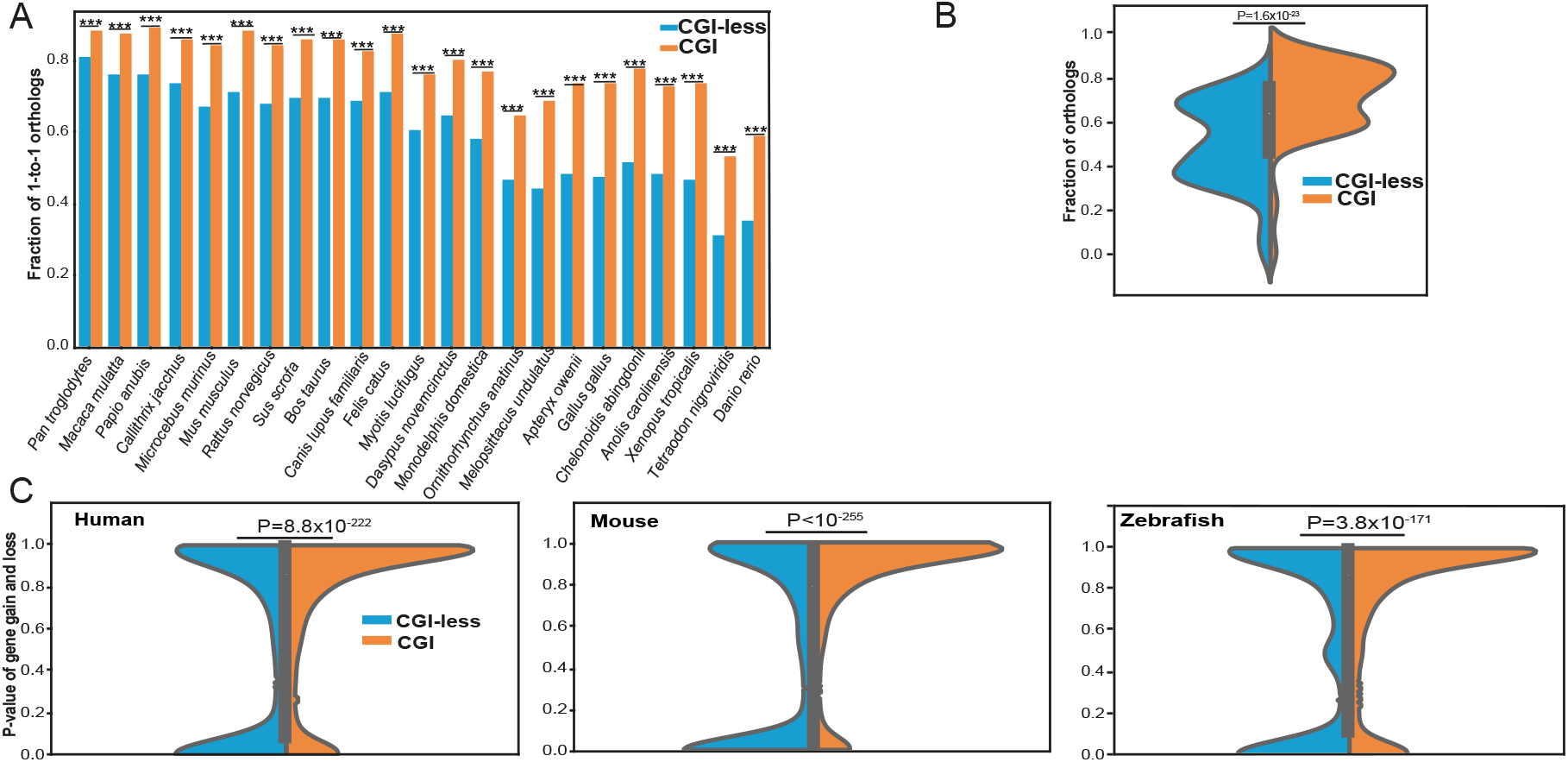
CGI and CGI-less genes duplication across species. (**A**) Fraction of 1-to-1 orthologs of human CGI and CGI-less genes (as defined using the conservative approach described in the main text) with a selected number of species, ordered by phylogenetic distance from human. In each pair of human -non-human species, CGI and CGI-less genes were compared using a chi-square test and corrected by FDR (*** -P<0.001). (**B**) Distributions of fractions of 1-to-1 orthologs across all 196 species that are annotated in ENSEMBL, in CGI versus CGI-less genes. Comparison between the distributions was preformed using a Mann-Whitney one-sided test. (**C**) Distributions of P-values of rates of gain and loss in CGI and CGI-less genes in human, mouse and zebrafish. Comparison between the distributions was performed using a Mann-Whitney one-sided test. See Fig S2 for additional species. In all species, CGI-less genes have a distribution enriched with low P-values, suggesting a higher rate of gene gain and loss in this group in comparison with CGI genes.

**Figure 2:**
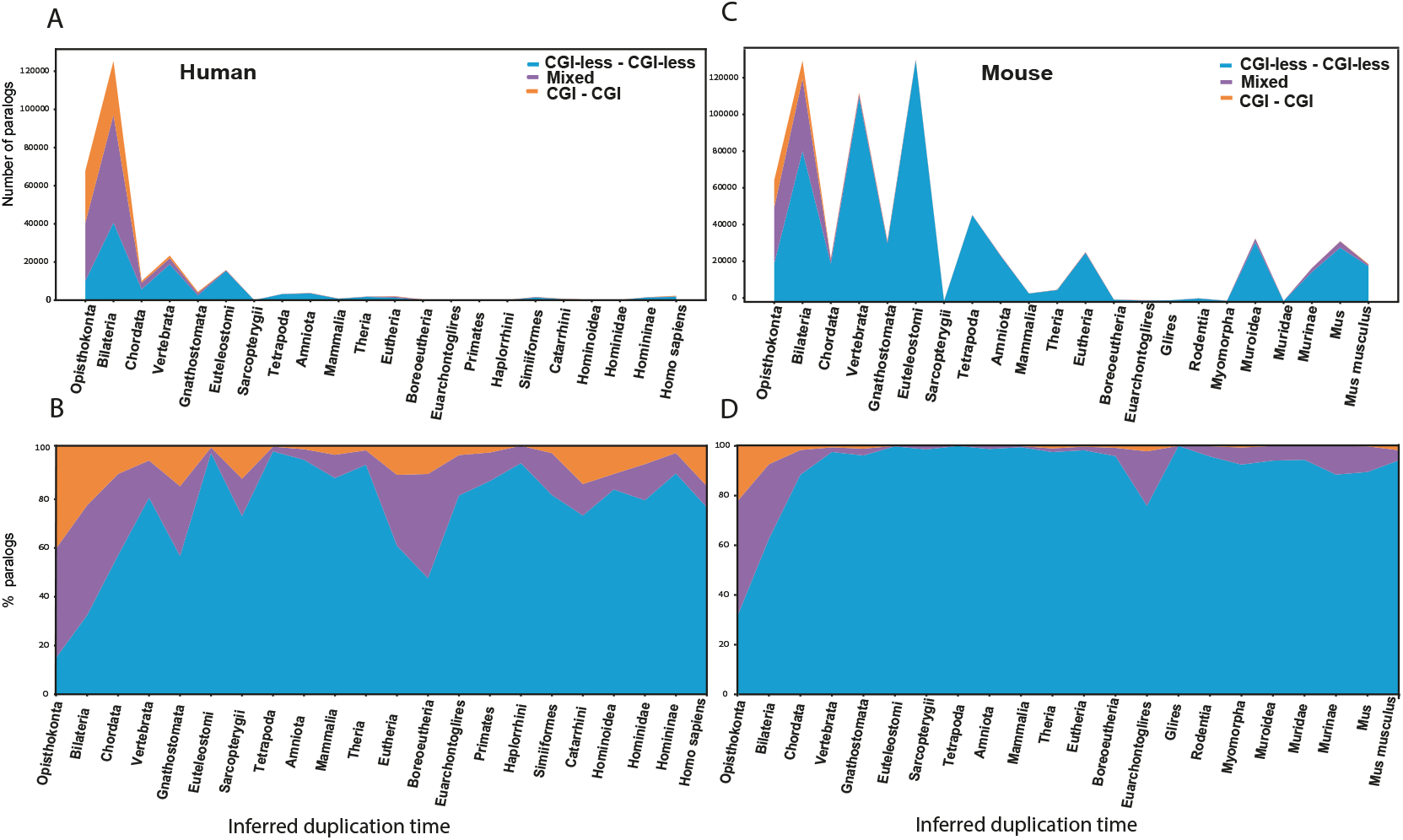
Evolutionary timeline of duplication events of CGI and CGI-less genes. (**A**) A timeline showing the total numbers of human paralogs. Each paralog is classified based on its inferred duplication time, ranging from Opisthokonta to *Homo Sapiens*. Paralogs are further partitioned based on their promoter classification: CGI -CGI paralogs (orange), CGI-less -CGI-less paralogs (blue) and Mixed (purple). CGI -CGI and Mixed pairs are skewed towards ancient times of duplication (permutation test, P<10^−5^). (**B**) A timeline showing the relative fractions of human paralogs at different duplication times, ranging from Opisthokonta to *Homo Sapiens*, where each point is split into CGI -CGI, CGI-less -CGI-less, and Mixed paralogs. (**C**) Similar to (A), with mouse paralogs. CGI -CGI and Mixed pairs are skewed towards ancient times of duplication (permutation test, P<10^−5^). (**D**) Similar to (B), with mouse paralogs.

These results suggest that CGI-less genes – that have fewer one-to-one orthologs across species in comparison with CGI genes, have undergone a higher rate of gene gain and loss events in the course of their evolution. Indeed, when looking at the rate of gene family expansion and contraction, as computed using the CAFE algorithm(30) and given in ENSEMBL Compara(31), we observe that CGI-less genes have a significantly faster rate of gene duplication and loss in comparison with CGI genes (**Fig 1C**).

We further tested this trend, of faster rate of gene gain and loss in CGI-less genes, in mouse genes where we used CGI annotations from the mouse genome, and observed the same phenomenon as seen in human genes (**Fig 1C** and **Supp Fig 3**). We next used experimentally verified data of non-methylated accessible regions (NMI regions, that correspond in function to CGIs) in genomes of less-studied species -chicken, anole lizard and zebrafish(29) to look at these trends of CGI versus CGI-less genes in analogous systems. In all cases, CGI-less genes show higher rates of gene gain and loss in comparison with CGI genes (**Fig 1C** and **Supp Fig 3**). Thus, CGI genes display lower rates of gene duplication and loss in the course of their evolution in comparison with CGI-less genes -an observation that holds in a diverse set of vertebrate species.

**Figure 3:**
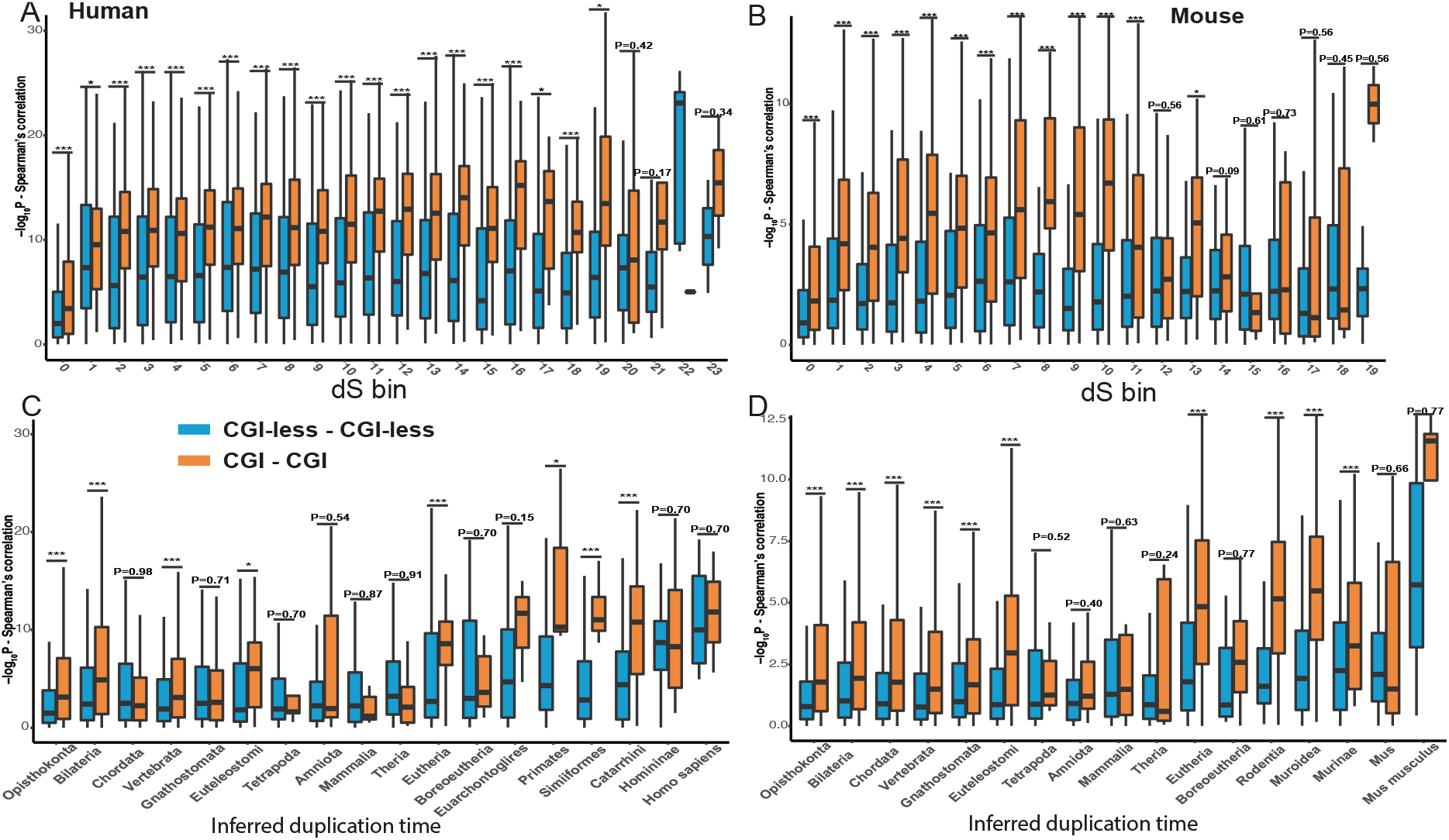
Transcriptional divergence in CGI and CGI-less paralogs. A timeline showing the transcriptional divergence of human (**A**,**C**) and mouse (**B**,**D**). Each paralog is classified based on its inferred duplication time. Paralogs are further partitioned to CGI-CGI and CGI-less -CGI-less pairs. Transcriptional divergence is computed based on Spearman’s correlation between the expression levels of the two paralogs across tissues and represented using the negative logarithm of the P-value of this correlation -a higher value represents a stronger agreement in expression and a higher correlation in transcription between the two paralogs. In **A-B**, paralogs are partitioned based on bins with similar values of synonymous substitutions (dS), with the lowest bin representing the most diverging paralog set (highest dS values). In **C-D**, the X-axis represnts the inferred duplication time based on tree topology. In each time point, the transcriptional diveregnce of CGI-CGI paralogs is compared with the CGI-less -CGI-less paralogs using a one-sided Mann-Whitney test. FDR-corrected P-values are shown. Inferred duplication times or dS bins with a small number of genes were removed in A-D. For clarity, only CGI-CGI and CGI-less -CGI-less paralogs are shown. The same plots including the Mixed paralogs are shown in Supp Fig 5. (***P < 0.001, **P < 0.01, *P < 0.05)

### Duplication events involving CGI genes are ancient and have mostly occurred before the emergence of vertebrates

To study the evolutionary patterns of gene duplication with respect to gene promoter architecture, we next focused on all pairs of paralogs in human and separately on all pairs in mouse. We divided paralog gene pairs into three categories, based on their promoter architectures: (1) **CGI paralogs** -where both paralogs are CGI genes, (2) **CGI-less paralogs** -where both paralogs are CGI-less genes, and (3) **Mixed paralog pairs** -where only one of the paralogs is a CGI gene. We note that the third group of Mixed paralogs is the least stable between the two methods of CGI gene classification, and we thus mostly focus on contrasting the two stabler groups -of CGI pairs with CGI-less pairs -in the following analyses.

We analyzed the inferred time of duplication of the three sets of paralogs in the course of evolution using either phylogenetically-based dating, based on gene tree topology from ENSEMBL Compara(31), or using the rate of synonymous substitutions, dS, between the two paralogs under the assumption it represents a molecular clock (see Methods). Both methods have been used in previous studies to estimate the time of duplication(10, 32, 33). We observe that those pairs that include CGI genes (either CGI paralogs or Mixed paralogs) almost entirely belong to ancient paralog categories in which the duplication events occurred before or around the time of vertebrate emergence. This is true for both human and mouse paralogs (**Fig 2A** and **2C** for human and mouse paralogs, respectively, and **Supp Fig 4** for dS-based analysis). Interestingly, the mouse genome includes a higher fraction of recently duplicated genes in comparison with human, including thousands of paralogs that their inferred last common ancestor is predicted to have existed in the murid ancestor. Regardless of their total number of recent duplications, in both genomes the overwhelming majority of these relatively recent events involves paralogs that lack CGI in the promoters of both genes (**Fig 2B** for human paralogs, P-value<10^−5^, permutation test; and **2D** for mouse paralogs; P-value <10^−5^, permutation test). For example, out of the total number of gene duplications that occurred after the last common ancestor of vertebrates, 87.1% and 97.1% are CGI-less paralog pairs in human and mouse, respectively.

**Figure 4:**
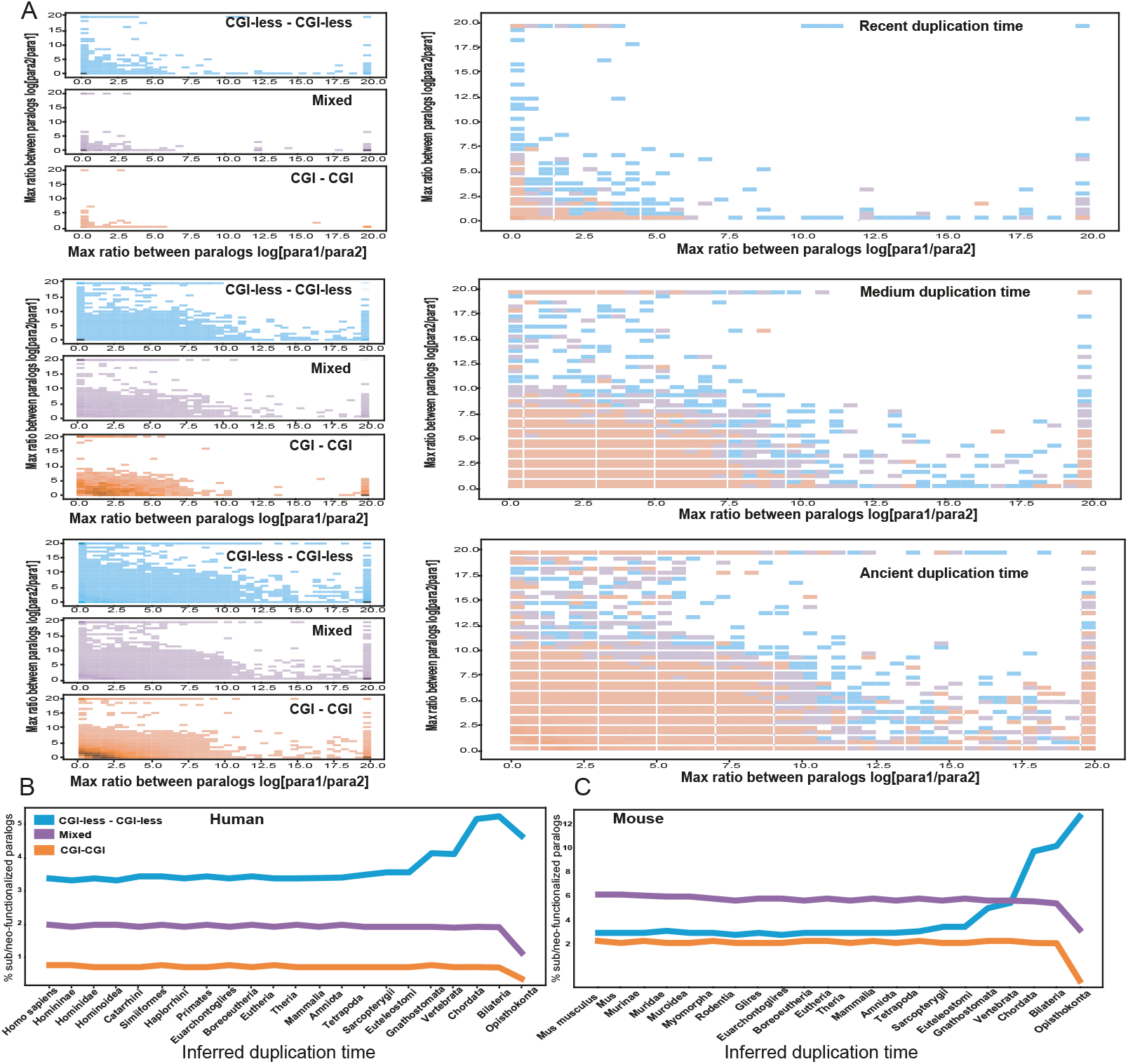
Transcriptional sub/neo-functionalization in CGI and CGI-less paralogs. (**A**) Heatmaps showing the log ratio of expression of human paralogs, where the X-axis represents the highest ratio between paralog1 and paralog2, and the Y-axis represents the highest ratio between paralog2 and paralog1 (chosen from the set of ratios of expression across tissues). Paralogs are split into recent, medium and ancient duplication times and to CGI-CGI, CGI-less – CGI-less and Mixed paralog classes. Left panels show the three classes separately, right panels show the aggregation of all classes together. Paralog pairs that appear in the top right corner tend to have more distinctive patterns of expression, where each of the two paralogs is strongly expressed in a different tissue. (**B-C**) The percentage of paralogs in human (B) and mouse (C) defined as “sub/neo-functionalized” based on low Spearman’s correlation and low Shannon entropy values out of the entire group of paralogs of their class. Paralogs are split based on their inferred duplication time and partitioned to CGI-CGI, CGI-less -CGI-less and Mixed paralog pairs. The fraction of paralogs that belongs to the “sub/neo-functionalized” group out of the total number of paralogs that are part of this particular class of paralogs is shown in each time point.

Thus, following the establishment of CGI as a major regulatory element in gene promoters during, or close to, the emergence of vertebrates(34), nearly all successful events of gene duplication and retention involved genes that are devoid of CGIs in their promoters. These observations are true for both human and mouse genes and are seen regardless of the method used to infer the time in which the duplication has occurred.

### Recent duplications of CGI-less genes are enriched with immune pathways and secreted proteins

We next asked whether gene duplication events at specific evolutionary periods, involving either CGI or CGI-less genes, are enriched in specific functions and pathways. Using GO term analysis(35) we looked for functional enrichment of non-redundant pairs of paralogs with specific promoter architecture (CGI or CGI-less) that have duplicated at particular evolutionary times (see Methods).

We observe that there are no shared functions or pathways in the majority of genes that are inferred to have duplicated at any given evolutionary period. This is expected as it suggests that gene duplication events occurring at specific evolutionary times are associated with multiple and unrelated pathways rather than with one major pathway. However, we did observe certain enriched functional categories for each of the paralog categories (CGI and CGI-less) in different duplication times -suggesting that particular pathways that require genes regulated by different promoter architectures have evolved through gene duplication during these evolutionary times (**Table 1**).

**Table 1:**
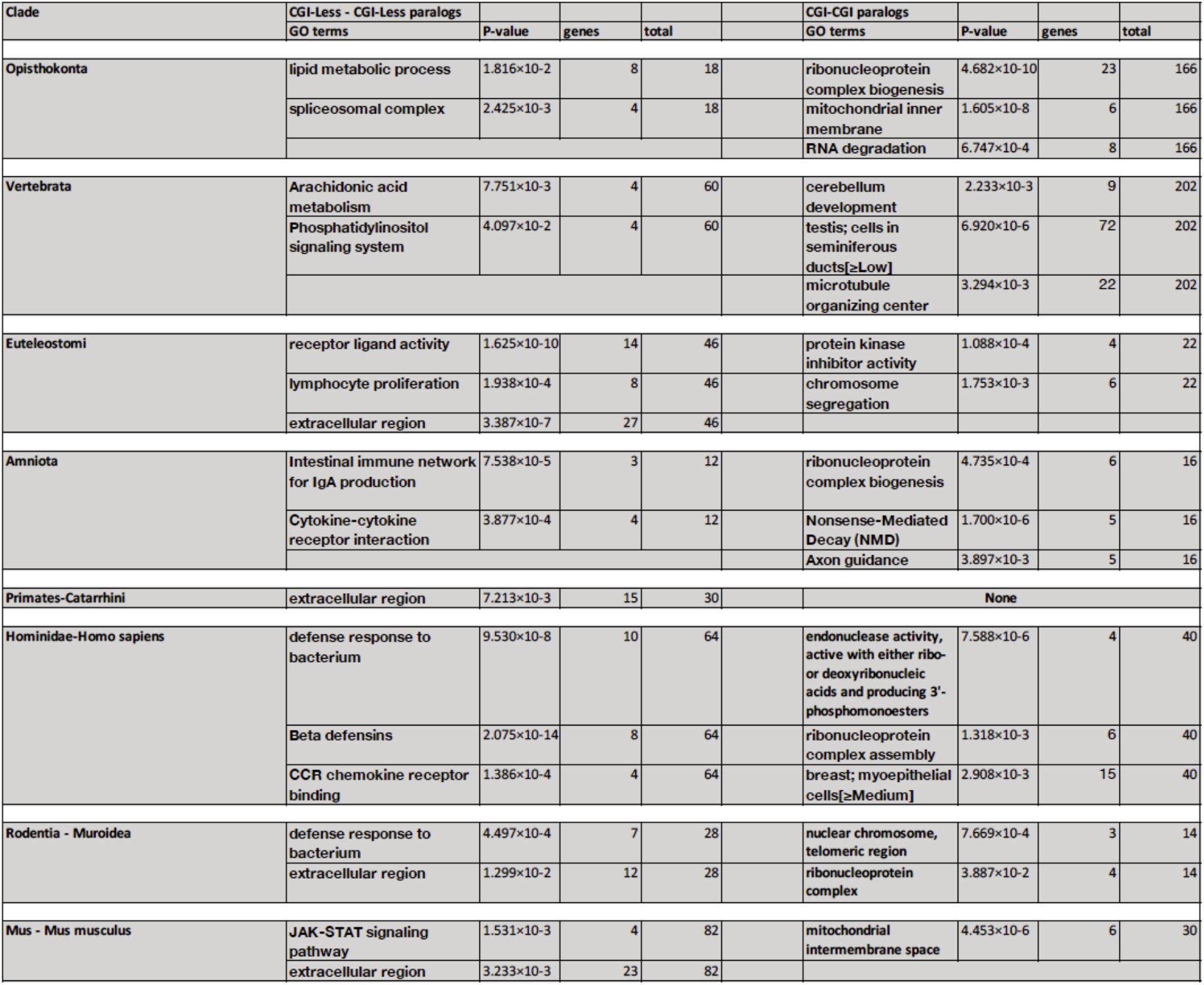
Go term enrichment of CGI and CGI-less paralogs in specific evolutionary periods. List and non-redundant enriched Go terms (with corrected P<0.05) in CGI-less – CGI-less paralogs thay have duplicated in ancient and recent evolutionary times.

| Clade | CGI-Less - CGI-Less paralogs |  |  |  | CGI-CGI paralogs |  |  |  |
| --- | --- | --- | --- | --- | --- | --- | --- | --- |
|  | GO terms | P-value | genes | total | GO terms | P-value | genes | total |
| Opisthokonta | lipid metabolic process | 1.816×10<sup>-2</sup> | 8 | 18 | ribonucleoprotein complex biogenesis | 4.682×10<sup>-10</sup> | 23 | 166 |
|  | spliceosomal complex | 2.425×10<sup>-3</sup> | 4 | 18 | mitochondrial inner membrane | 1.605×10<sup>-8</sup> | 6 | 166 |
|  |  |  |  |  | RNA degradation | 6.747×10<sup>-4</sup> | 8 | 166 |
| Vertebrata | Arachidonic acid metabolism | 7.751×10<sup>-3</sup> | 4 | 60 | cerebellum development | 2.233×10<sup>-3</sup> | 9 | 202 |
|  | Phosphatidylinositol signaling system | 4.097×10<sup>-2</sup> | 4 | 60 | testis; cells in seminiferous ducts[≥Low] | 6.920×10<sup>-6</sup> | 72 | 202 |
|  |  |  |  |  | microtubule organizing center | 3.294×10<sup>-3</sup> | 22 | 202 |
| Euteleostomi | receptor ligand activity | 1.625×10<sup>-10</sup> | 14 | 46 | protein kinase inhibitor activity | 1.088×10<sup>-4</sup> | 4 | 22 |
|  | lymphocyte proliferation | 1.938×10<sup>-4</sup> | 8 | 46 | chromosome segregation | 1.753×10<sup>-3</sup> | 6 | 22 |
|  | extracellular region | 3.387×10<sup>-7</sup> | 27 | 46 |  |  |  |  |
| Amniota | Intestinal immune network for IgA production | 7.538×10<sup>-5</sup> | 3 | 12 | ribonucleoprotein complex biogenesis | 4.735×10<sup>-4</sup> | 6 | 16 |
|  | Cytokine-cytokine receptor interaction | 3.877×10<sup>-4</sup> | 4 | 12 | Nonsense-Mediated Decay (NMD) | 1.700×10<sup>-6</sup> | 5 | 16 |
|  |  |  |  |  | Axon guidance | 3.897×10<sup>-3</sup> | 5 | 16 |
| Primates-Catarrhini | extracellular region | 7.213×10<sup>-3</sup> | 15 | 30 | None |  |  |  |
| Hominidae-Homo sapiens | defense response to bacterium | 9.530×10<sup>-8</sup> | 10 | 64 | endonuclease activity, active with either ribo- or deoxyribonucleic acids and producing 3'-phosphomonoesters | 7.588×10<sup>-6</sup> | 4 | 40 |
|  | Beta defensins | 2.075×10<sup>-14</sup> | 8 | 64 | ribonucleoprotein complex assembly | 1.318×10<sup>-3</sup> | 6 | 40 |
|  | CCR chemokine receptor binding | 1.386×10<sup>-4</sup> | 4 | 64 | breast; myoepithelial cells[≥Medium] | 2.908×10<sup>-3</sup> | 15 | 40 |
| Rodentia - Muroidea | defense response to bacterium | 4.497×10<sup>-4</sup> | 7 | 28 | nuclear chromosome, telomeric region | 7.669×10<sup>-4</sup> | 3 | 14 |
|  | extracellular region | 1.299×10<sup>-2</sup> | 12 | 28 | ribonucleoprotein complex | 3.887×10<sup>-2</sup> | 4 | 14 |
| Mus - Mus musculus | JAK-STAT signaling pathway | 1.531×10<sup>-3</sup> | 4 | 82 | mitochondrial intermembrane space | 4.453×10<sup>-6</sup> | 6 | 30 |
|  | extracellular region | 3.233×10<sup>-3</sup> | 23 | 82 |  |  |  |  |

For example, CGI duplications that are inferred to have occurred at the root of the vertebrate clade are associated with genes related to tissue-specific function and identity, including cerebellum and testis (FDR-corrected P-value= 2.2×10^−3^ and 6.9×10^−6^, respectively). Interestingly, CGI-less duplications that are inferred to have occurred after the Euteleostomi last common ancestor are enriched with various immune-related pathways as well as extracellular and secreted proteins. This enrichment of immune and secreted proteins is also noticeable in very recent duplications, defined as those that occurred following the split between primates and rodents, and is observed in both lineages that lead to human and mouse (**Table 1**). The enrichment of recent gene duplications in immune functions was also observed in previous comparative analyses of primate(36) and rodent(37) genomes, and here we show that these duplication are associated with CGI-less genes.

### CGI paralogs display lower transcriptional divergence in comparison with CGI-less paralogs

Previous studies suggested that genes lacking CGI in their promoters display greater plasticity in their expression – transcriptionally varying to a greater extent than CGI genes across tissues, individual cells and conditions(27, 28, 38). Furthermore, orthologous CGI-less genes diverge more in their expression across species with respect to CGI genes(27, 28). This raises the question whether CGI-less paralogs have greater transcriptional divergence between the two paralogs, in comparison with CGI paralog pairs. In other words, previous work showed that promoter architecture is linked to changes in gene expression of the same gene in different conditions. Here we ask whether promoter architecture of paralog genes, in the form of presence or absence of CGIs, is linked to differences in expression between the two paralogs, which can have important consequences for paralog sub-and neo-functionalization.

To test this, we looked at the differences between paralogs in their expression across a large set of tissues in human and in mouse. We used the GTEx data, which includes transcriptomics data from numerous tissues from a large number of human donors(39) and the smaller mouse transcriptomics BodyMap dataset(40) to compare paralog expression in human and mouse tissues, respectively. We computed the Spearman’s correlation of each paralog pair’s expression across the tissues: low Spearman’s correlation denotes transcriptional divergence between the paralogs, while high Spearman’s correlation suggests stronger similarity in expression between them. We split the pairs into different paralog categories (CGI, Mixed and CGI-less). We further partitioned the paralogs based on their inferred duplication time (**Fig 3 and Supp Fig 5)**. We partitioned them using both methods mentioned above to infer the time of duplication – dS values between paralogs and dating based on tree topology. We use both methods due to noise in the transcriptional data.

**Figure 5:**
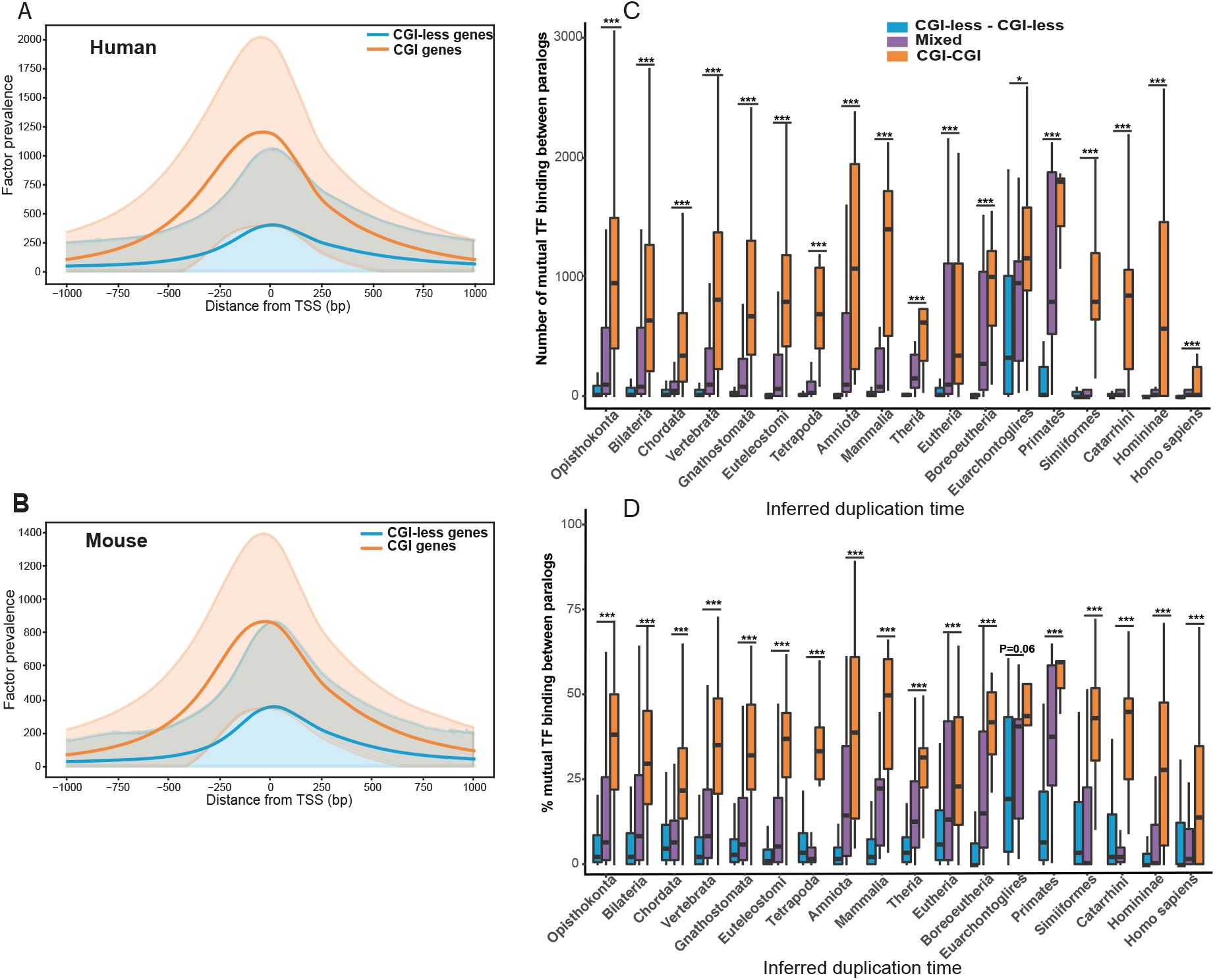
TF-binding in promoters of CGI and CGI-less genes and paralogs. (**A**) A TSS-relative histogram of TF ChIP-seq peaks from the Cistrome dataset for CGI and CGI-less genes. The cumulative number of TF ChIP-seq peaks from the Cistrome dataset that intersect with promoter regions of CGI and CGI-less human genes are shown. The shaded regions represent one standard deviation from the mean. (**B**) Similar to (A), with mouse genes and with the mouse ChIP-seq data from Cistrome. Both human and mouse CGI genes have a greater number of TF binding to their promoter regions in comparison with CGl-less genes (P-value<1×10-^307^, t-test). (**C**) Total number of shared TFs that bind to the promoters of both paralogs for human CGl-CGl, CGl-less -CGl-less and Mixed pairs. Paralogs are partitioned based on their inferred duplication time. ln each time point, the distributions of shared TF numbers are compared between CGI-CGI paralogs and CGI-less – CGI-less paralogs using a one-sided Mann-Whitney test. P-values are corrected by FDR. (**D**) Relative fraction of shared TF binding to the promoters of both paralogs in human CGI-CGI, CGl-less -CGl-less and Mixed paralog pairs. ln each time point, the distributions of CGl-CGl paralogs and CGl-less -CGl-less paralogs are compared using a one-sided Mann-Whitney test. P-values are corrected by FDR. In nearly all time points, the total number (C) and the relative fraction (D) of the shared TFs between paralogs are significantly larger in CGl-CGl in comparison with CGl-less -CGl-less paralogs. Analogous analyses in mouse give similar results (Supp Fig 9). (***P < 0.001, **P < 0.01, *P < 0.05)

In general, we observe a trend where younger duplicates are more similar to each other in their transcriptional profiles, as inferred from the higher Spearman’s correlation values between these paralogs, whereas ancient duplicates are more divergent (**Fig 3**). For example, paralogs with the largest dS values between the paralog pairs – those that are likely to have duplicated in ancient times and belong to the first dS bin in our data – have a median value of Spearman’s ρ of 0.37 and 0.32 in human and mouse, respectively. In contrast, the corresponding values in paralogs with smaller dS values (all remaining genes that do not belong to the first bin in our data) are ρ= 0.71 and 0.42 in human and mouse, respectively. This greater transcriptional divergence and lower correlation between ancient paralogs is expected, given the longer time since duplication, and is in agreement with previous observations(33, 41).

Importantly, when comparing paralogs that are inferred to have duplicated at similar times, we observe that in the majority of the cases CGI paralogs are transcriptionally more similar to each other in comparison with CGI-less paralogs (**Fig 3**). Conversely, CGI-less paralogs show greater transcriptional divergence in comparison with CGI paralogs. This finding is observed in both human and mouse, suggesting a greater capacity of CGI-less genes to transcriptionally diverge and to potentially give rise to neo-or sub-functionalization between CGI-less paralogs. We note that this trend is not observed in recently duplicated genes, which may reflect the requirement for longer time since duplication to diverge(32). Alternatively, this comparison may suffer from the small number of recently duplicated CGI-CGI paralogs, reducing the statistical strength of this analysis.

### CGI paralogs display lower rates of neo-and sub-functionalization than CGI-less paralogs

Further to the above findings of greater transcriptional divergence between CGI-less paralog genes, we next asked whether CGI-less paralogs also show an increased tendency to sub-or neo-functionalize. While gene duplication is thought to be an important mechanism for evolving new functions, recent work suggested that the process by which gene duplicates transcriptionally evolve to sub-or neo-functionalize is often slow, and is rarely observed in relatively recent duplication events(32). To investigate transcriptional sub-and neo-functionalization in paralogs with different promoter architectures, we used the GTEx data to look at how contrasting the paralog gene expression is between the two paralogs across human tissues. This was done by finding the two most extreme tissues where the two paralogs are most highly and lowly expressed in comparison to each other. For this, we computed the ratio of expression between the two paralogs in any given tissue (log[paralog1/paralog2]), taking care of cases where one or both paralogs are lowly expressed (see Methods). For each pair, we found the tissue where the ratio is the highest for the first paralog and the tissue where this ratio is the lowest (and thus highest for the second paralog). We then plotted these two values against each other (such that one axis represents the highest ratio of paralog1/paralog2, and the other axis represents the highest ratio of paralog2/paralog1). We split the paralog pairs based on their inferred time of duplication into three major evolutionary periods – ancient, middle and recent times, and within each time category, between the three promoter classes (see **Fig 4A** and Methods for details). As expected, more contrasting values between the paralogs are seen in older paralog pairs, given their longer time of divergence since duplication. Importantly, we observe that in all three evolutionary periods, there is a greater tendency of CGI-less paralogs to have larger differences between the two paralogs. Similar patterns are observed when looking at the mouse paralogs using the BodyMap transcriptomics dataset (**Supp Fig 6**). This points to a greater capacity of CGI-less paralogs to sub-or neo-functionalize and to be uniquely expressed in different tissues, in comparison with paralogs with CGI promoters.

To test this further, we have adopted a previous approach to define pairs as being sub-or neo-functionalized, where Shannon entropy is computed for each gene’s expression values across different tissues to indicate the level of tissue-specificity in expression(42). Lower Shannon entropy denotes a stronger tendency for tissue-specific expression, while higher entropy implies a more uniform cross-tissue expression. Using low Shannon entropy values combined with the low Spearman’s correlation values of expression of the two paralogs, we can identify the set of paralog pairs that are most distinctively expressed across tissues among all paralog pairs, and are thus more likely to be sub-or neo-functionalized (see Methods). Using this approach, and in agreement with previous results(32), we find that only a small fraction of recent duplication events belongs to these “distinctively expressed pairs”, while a higher fraction of ancient pairs is part of this category in both human and mouse (**Fig 4B**). When comparing the fraction of the three classes of paralogs (CGI, CGI-less and Mixed paralogs) belonging to the distinctively expressed pairs, we observed that a larger fraction of human CGI-less pairs is distinctively expressed with respect to their total numbers (**Fig 4B**). When performing the same analysis with the mouse data, we observed that the same is true in ancient paralogs, where CGI-less paralogs dominate the distinctive expression category of paralogs. However, in more recent duplications in the mouse lineage – from the ancestor of vertebrates – the Mixed pairs have higher fractions of distinctively expressed paralogs than the CGI-less pairs (**Fig 4C**). This partial discrepancy between human and mouse may represent a truly biological difference in CGI regulation of genes between human and mouse, but it can also originate in the more limited expression data available for mouse.

Taken all the above results together, CGI-less pairs display greater transcriptional divergence in comparison with CGI paralogs, and this may have contributed to a greater number of CGI-less paralogs that are sub-or neo-functionalized in their expression across tissues.

### CGI paralogs share greater similarity of TF binding patterns between their promoters than CGI-less paralogs

The higher transcriptional divergence observed in CGI-less paralogs in comparison with CGI paralogs may point to larger differences in *cis*-regulation between CGI-less paralogous genes. To test this notion, we compared transcription factor (**TF**) binding in promoter regions of paralogs, using Cistrome -a large dataset of ChIP-seq data that assembles numerous TF-ChIP studies in both human and mouse in a diverse set of cells and tissues(43). Following exclusion of several general TFs and insulators (see Methods), we overlapped each gene’s promoter region with peaks of various TF-ChIP-seq data, yielding the set of TFs that are experimentally shown to bind the proximal promoter region of each human and mouse gene.

With this data, we first compared the total numbers of TFs that bind to the promoters of CGI and CGI-less genes across cells and tissues, and observed that a significantly larger total number of binding events is recorded for CGI genes (**Fig 5A-B**, P-value<1×10^−307^, t-test). We also note that there are differences in the relative occupancy of TFs along the promoter region, where a wider region upstream of the TSS is occupied by TFs in CGI genes and a narrower region of occupancy is observed in CGI-less genes (**Supp Fig 7** P-value<1×10^−130^, t-test). Finally, the pattern of promoter sequence conservation near the TSS differs between CGI and CGI-less genes (see **Supp Figs 8** and Methods for details). Thus, CGI and CGI-less genes significantly differ in the profiles of TF bindings to their proximate promoters, in agreement with previous findings(25, 27, 44).

We next compared the sets of TFs that bind to promoters of paralog pairs, to estimate the similarity in TF binding events between paralogs. We observed that CGI paralogs have significantly higher total numbers of shared TFs that bind to both paralog promoters in comparison with CGI-less paralogs (**Fig 5C** and **Supp Fig 9A**). Furthermore, even after normalizing for the higher total number of TFs that bind to CGI genes, which we discussed above, we observe that a significantly higher fraction of the TF binding events is shared between CGI paralogs than in CGI-less paralogs (**Fig 5D** and **Supp Fig 9B**). Thus, both in absolute and in relative numbers, there is a greater similarity between CGI paralogs, with respect to TF binding to their promoters, than between CGI-less paralogs. These findings, which were observed for both human and mouse genes, suggest that at least part of the greater transcriptional divergence in CGI-less paralogs can be attributed to the larger differences between the two paralogs in the binding of TFs to their proximal promoters. In contrast, CGI-CGI paralogs share a higher fraction of TFs that bind to both promoters, which likely supports the observed lower transcriptional divergence in these paralogs.

## Discussion

Gene duplication creates novel genetic material that has the potential to facilitate the emergence of new functions. However, it also entails maintaining gene dosage between the two duplicates as well as necessitating the evolution of distinctive functions for the successful fixation of the majority of paralogs. Here, we analyzed the links between gene duplication and retention and gene promoter architecture, as manifested in the presence or absence of CGIs – a major regulatory element in vertebrate genomes. We first observed that nearly all duplication events of CGI genes are ancient and have likely occurred before the establishment of CGI as an important promoter element, close to the origin of vertebrates(34). In contrast, CGI-less genes dominate successful events of gene duplication and retention in more recent times. These results corroborate previous studies that highlighted the differences in function and expression between genes that duplicated as part of ancient whole genome duplication events (ohnologs) versus tandem duplication events(8, 45). While ohnologs were found to be enriched for ubiquitously expressed genes, paralogs from small-scale duplications were more tissue-restricted(45) – in agreement with known characteristics of CGI and CGI-less genes, respectively. Thus, many cases of CGI gene duplication and retention are associated with ancient whole genome duplication and have supported the increase in copy numbers of ubiquitously expressed genes.

More recent duplications, occurring through small-scale duplication, involve CGI-less genes that tend to be more tissue-or condition-specific genes, such as those that function in immune and extracellular contexts. Both immune and secretion classes include many induced genes whose expression significantly varies between conditions, likely reflecting the requirement for having a promoter that lacks CGI to support this type of induced or plastic transcriptional regulation. The enrichment we found, of immune gene duplication in recent times, suggests that gene duplication may play an important role in mammalian adaptation to pathogens, in addition to the known rapid coding sequence evolution that is observed in many immune genes(46, 47). Furthermore, this recent duplication of immune genes mostly happens in CGI-less promoter genes, in agreement with results that suggest that the most rapidly evolving class of immune genes lack CGI in their promoters(27, 28).

A major factor that impacts gene’s transcriptional divergence is the regulatory structure of the proximate promoters. Previous studies have repeatedly pointed to a greater transcriptional plasticity of genes that lack CGI in their promoters(27, 28, 38, 48). In contrast, lower variability and greater homogeneity in expression was observed in CGI-genes. Furthermore, orthologous CGI-genes are more conserved in their expression across species, giving rise to suggestions that CGI promoters are more robust to mutations(28), although the molecular mechanisms behind this remain largely unclear. We here show that this is also true for paralogs: CGI-less paralogs exhibit greater transcriptional divergence between the paired genes across tissues, as well as a greater tendency to sub-or neo-functionalize. This higher transcriptional divergence between paralog pairs of CGI-less genes is also mirrored in larger differences in the set of TFs that bind the promoters of CGI-less gene pairs.

It is likely that the observed preferential retention of duplicated CGI-less genes in recent times – after the last common ancestor of vertebrates – reflects a combination of several factors: (1) Many of the cellular pathways and functions that have undergone major alterations in recent evolutionary times involve genes that are expressed in a specific set of conditions or tissues, such as immune-related genes, and this mode of expression is supported by promoters lacking CGIs; (2)CGI-genes are often highly expressed across many tissues, and their duplication may involve complex dosage compensation processes that are easier to resolve in CGI-less genes whose transcription seems more plastic and amenable to changes; (3) Functional CGI promoters may be harder to evolve, requiring an elongated CGI region, unlike CGI-less promoters that have a shorter effective proximal promoter with a few TF binding modules (as observed in Fig **5A-B** and **Supp Figs 7-8**). The latter two factors, dosage compensation and promoter evolvability, are more important in small-scale duplications, the main mode of duplication in recent duplication events.

Our results thus provide a detailed chronology of paralog accumulation before and during vertebrate evolution with respect to gene promoter architecture and their subsequent divergence in transcription and in *cis*-regulation. They further demonstrate how promoter features influence transcriptional evolvability and, in turn, the successful retention of new genes, enabling evolutionary innovation.

## Methods

### Gene and genome annotation

We downloaded gene annotations, including orthology and paralogy assignments, from ENSEMBL version 98, which corresponds to GRCh38, GRCm38, GRCg6a, AnoCar2.0 and GRCz11 genome assemblies for Human, Mouse, Chicken, Anole lizard, and Zebrafish, respectively. We have removed genes that are not coding for proteins or whose transcripts are not known, and kept only the primary assembly genes. Similarly, for pairs of paralogous genes, we have only included pairs of genes where both genes are coding, resulting in a total of 133,328 pairs of paralogs in humans and 356,568 pairs in mouse.

We separate paralogs based on their inferred time of duplication based on two different and commonly used methods(10, 32, 33): (1) Inferred duplication time based on ENSEMBL tree and provided by ENSEMBL Compara(31), and (2) based on the synonymous substitution rate – dS, between the two paralogs (where higher dS values imply longer time since duplication(32, 49, 50)). For the latter method, we binned paralog pairs into 24 bins: Paralogs that significantly diverge in sequence (i.e. with high dS values, above 2), were binned into a single bin, which likely includes many ancient category of paralogs. This resulted in the first bin being much larger than the other 23 bins. In addition, all zero-value dS paralogs were binned into the 24th bin, which is likely enriched with recent duplications. This resulted in all median bins (2nd-23rd) to have a similar size, while the first and the last bins are larger than the rest.

In the Results section, we show most of the analyses using the first method (dating based on ENSEMBL tree topology). In the case of the relative abundance of different paralog classes in the course of evolution (**Fig 2** and **Supp Fig 4**) and when in transcriptional divergence analysis (**Fig 3** and **Supp Fig 5**) we show the results using both approaches.

### CGI and Promoter classification

For the human genome, CpG Island (CGI) annotations were downloaded from ENSEMBL as well as from UCSC Genome browser(51) [https://genome.ucsc.edu/cgi-bin/hgTables] (Assembly 2013/12 hg38). These annotations gave nearly identical results (see **Supp Fig 1**). We show all following analyses using the UCSC Genome browser annotations. Since CGI predictions in species other than human were shown to diverge from experimental data of non-methylated regions(29), we used ENSEMBL CGI annotations for mouse (as ENSEMBL also includes experimental data). For all other species – chicken, lizard and fish, which lack extensive data of their CGIs, we used experimental data of non-methylated islands (NMI) from liver tissues(29). For two species, we used liftover(52) to get the coordinates of NMI regions in more recent genome versions: Danio rerio (DanRer11 lifted over from DanRer7) and Anolis carolinensis (AnoCar2). For Gallus gallus, we repeated the peak calling procedure with the chicken liver data from GSE43512 accession(29), using galGal6.

We defined CGI genes – genes harboring CGIs in their promoters – as genes that at least 50% of the region spanning from 300bp upstream of the TSS and 100bp downstream it overlap with annotated CGIs (or NMIs), as previously done(27, 28). All other genes were defined as CGI-less genes. Transcription Start Site (TSS) coordinates were taken from the canonical transcript (defined as the longest transcript among the gene’s transcripts, which has the lowest TSL (transcript support level), or the longest transcript if no TSL transcripts exists) for each gene. We also used an alternative approach to define CGI-genes (a more ‘relaxed’ approach), using a minimum 1bp overlap of annotated CGIs with the region of 1,000bp both up-and downstream of the TSS, following previous work(29). All following analyses are shown using the first approach and the conclusions hold regardless of the method used. For comparison, we also show the results of Main **Fig 1** using the second approach, in **Supp Figs 2-3**.

These CGI definitions for each gene, enabled us to define three groups of paralogs, based on their CGI status: (1) CGI paralogs -where both paralogs are CGI genes, (2) CGI-less paralogs -where both paralogs are CGI-less genes and (3) Mixed paralog pairs -where only one of the genes is a CGI gene, denoted respectively as “CGI-CGI”, “CGI-less – CGI-less” and “Mixed” pairs throughout the article. The third group of Mixed paralogs was the least stable across the two methods used to classify CGI genes, and thus most analyses focus on contrasting the two stabler groups of CGI pairs with CGI-less pairs.

### Functional enrichment analysis

To study enrichment for cellular functions and pathways in genes that are inferred to have undergone duplication in different evolutionary time-points, which have a particular promoter architecture (CGI-CGI versus CGI-less – CGI-less paralog pairs), we used only pairs of paralogs that do not belong to large gene families. This was done to avoid gene families with numerous paralogs whose inferred time of duplication may be obscured by the complex relationship among various paralog members in the family. We thus focused on a non-redundant set of paralog pairs, where each paralog pair has no other paralog. We note that the majority of paralog pairs belong to this category, both in human and in mouse. We used gProfiler(35) to find enriched pathways within each subset of paralogs that (1) have a particular promoter architecture and (2) are inferred to have duplicated at the same evolutionary time (e.g. the set of all CGI-CGI paralogs that are inferred to have duplicated with their last common ancestor found in the vertebrate ancestor). This enrichment was done with default settings (i.e., against the background of all genes). The results of the significantly enriched functions (FDR-corrected P-values<0.05) are shown in Table 1. In this analysis we only considered pathways and functions that are associated with more than two genes and removed statistically-enriched pathways with fewer genes, to avoid cases where the enrichment is based only on one pair of paralogs.

### Transcriptional divergence analysis

To study the expression patterns of paralogs across a large set of tissues we used RNA-seq data from the Genotype-Tissue Expression (GTEx) project, version 8(39) – a large transcriptomics dataset with gene expression across different tissues from a large number of human individuals. We filtered out all the pseudoautosomal expression records, along with non-primary tissues (cultured cells, EBV-transformed lymphocytes and CML). For all expression-based analyses, we followed the same filtering used in Lan et al.(32), by removing any genes with total expression below 5 TPMs across all tissues as well as those whose expression levels are below 0.5 TPM in every single tissue. Finally, for all the transcriptomics-based analyses, we only used paralog pairs with less than 80% sequence similarity between the two paralogs, excluding highly-similar pairs. We estimated the similarity in expression of each pair of paralogs across tissues, by computing the Spearman’s rank correlation coefficient, ρ, between the two genes. For this, we used the median TPM values of the two paralogs across the 51 GTEx tissues. High correlation value denotes high similarity in expression profiles between the two paralogs, whereas low correlation indicates discrepancy in expression between the two genes across different tissues. Following this, we computed the distribution of ρ values in each class of paralogs (CGI-CGI, CGI-less – CGI-less and Mixed), splitting them further based on the inferred time in which the duplication has occurred (Fig 3). We used both ENSEMBL tree topolgy dating and dS bins to classify the paralogs by their inferred duplication time, and in the case of the first method removed small groups from the analysis. To compare the distributions of CGI-CGI with CGI-less – CGI-less paralogs, we used a one-sided Mann-Whitney test, followed by an FDR-correction.

We used a similar approach with the mouse BodyMap dataset(40), where mean expression of each gene across 17 tissues, split by sex into 33 different tissues, was used for computing the Spearman’s rank correlation scores. TPM values were quantified by downloading raw fastq files from PRJNA375882 accession, using fasterq-dump from the SRAtools package (https://github.com/ncbi/sra-tools), followed by quantification using Salmon, version 1.4.0(53), in mapping mode on an indexed transcriptome of mouse from ENSMEBL version 98.

### Sub/Neo-functionalization analysis using relative expression levels

To quantify the tendency of any given pair of paralogs to sub-or neo-functionalize, we searched for the two tissues where the ratio of expression between the two paralogs is most extreme. This was done such that in the first tissue the first paralog is most highly expressed in comparison with the second paralog’s expression, whereas in the second tissue the opposite is true (the second paralog is highly expressed with respect to the first paralog). We took care of cases, where one or both paralogs are lowly expressed in the following matter:

‘Tissue similarity’ was computed by applying exp_diff function to the medians of each of tissue expression in each of the genes, as follows:

def exp_diff(x,y):

RANGE=20

if x<y:

return -exp_diff(y,x) if x<1:

return 0

elif x>2 and y>x/exp(RANGE):

return min(RANGE, log(x/y))

elif x>2 and y<=x/exp(RANGE):

return RANGE

elif y>x/exp(RANGE)

return log(x/y)*(x-1)

else:

return RANGE*(x-1)

Where x and y are expression levels (TPMs) of the two paralog genes in a given tissue.

The above procedure took care of the result being (1) symmetric, (2) roughly logarithmic, (3) bounded, (4) smooth and (5) differentiable almost everywhere. It also took care of gene expression level, suppressing noise from fluctuations of lowly-expressed genes (defined in this analysis as genes with <1 TPM in the tested tissue, and bounding the range when such genes pair with well-expressed genes. We carried out this procedure on all filtered human and mouse paralog pairs that we have analyzed their expression using the GTEx and the BodyMap, respectively (as described above).

We next partitioned the paralogs based on their inferred time of duplication, into three groups -“ancient”, “middle” and “recent”, as follows: For human -duplications occurring in the Primates clade and younger duplications were defined as “recent”. For mouse -duplications occurring between the ancestor of Mus musculus to the ancestor of Glires were defined as recent. For both human and mouse, paralogs that their last ancestor before duplication is inferred to be at the Opisthokonta or Bilateria ancestral nodes were defined as “ancient” paralogs, and all paralogs inferred to duplicate between the “ancient” and the “recent” periods were defined as “middle”.

Paralogs belonging to each of these three periods were plotted separately on a heatmap. For each paralog pair, we took the two values of relative expression from the two extreme tissues (the logarithm of expression between paralog1 and paralog2 in the tissue where paralog1 is most highly expressed with respect to paralog2, and the opposite for the second tissue, as explained above). These heatmaps (**Fig 4A** and **Supp Fig 6**) convey for each class of paralogs and in each evolutionary period the contrast in expression between paralog pairs, where more contrasting values tend to be higher on both axes. Thus, paralogs pairs that are more contrasting in their expression (in at least two extreme tissues) are located towards the top-right of each heatmap.

### Sub/Neo-functionalization analysis using an intersection of Shannon entropy measurement and Spearman’s rank correlation

We implemented an additional approach to detect a subset of paralogs that have neo-or sub-functionalize in their transcription across tissues: In this second procedure, we first computed Shannon entropy for each gene’s expression across different tissues to indicate the level of tissue-specificity in expression(42). Lower Shannon entropy denotes stronger tendency for tissue-specific expression, while higher entropy implies a more uniform cross-tissue expression. Paralog pairs may have sub-/neo-functionalized if both genes are expressed in a tissue-specific manner, with low correlation in expression between the two paralogs – i.e., if the two paralogs tend to be highly expressed, each in a different set of tissues.

To find this set of tissue-specific and lowly-correlating paralogs, we combined measurements of (1) Shannon entropy and (2) Spearman correlation of the paralog expression across tissues, as measured in the GTEx or BodyMap data. We can define sub-or neo-functionalized paralog pairs as those pairs where: (1) each of the two genes belongs to the lowest quartile of Shannon entropy values, as well as (2) the two genes having a low Spearman’s correlation value across tissues, belonging to the lowest quartile of Spearman’s correlation values. This method thus enriches for paralog pairs expressed differently across tissues with respect to each other, and in a tissue-specific manner.

We next plotted the fraction of the paralogs that belong to this class of sub/neo-functionalized pairs, out of the total group of that paralogs. We partitioned the paralogs based on their promoter architecture and based on their inferred time of duplication, as we did in Figs 2-3. We repeated this analysis for both human and mouse paralogs (**Fig 4B-C**).

### Transcription factor binding analysis

ChIP-seq data (in the format of narrowPeak bed files) of a large set of human and mouse transcription factors binding experiments were downloaded from the Cistrome database(43, 54). From this data we excluded a few general factors and insulators: CTCF, RAD21, REST, EP300, and RNA Polymerase (POLR), and removed TFs with no binding recorded. With the filtered TF-binding data, we counted for each of the human and mouse genes which and how many TFs intersect in their binding region with the gene’s promoter area. This was done by testing the overlap of at least one base pair between the promoter region and the peaks in the narrowPeak bed files. Following this, we first examined the total number of TF binding events for each CGI and CGI-less genes (**Fig 5A-B**) in a region of 2,000bp up-and downstream of the TSS of each gene. We also plotted a normalized histogram of this in **Supp Fig 7**.

We next compared the similarity of TF binding to promoters of pairs of paralogs, partitioned based on their promoter classes. This was done by looking at (1) the total number of shared TF bindings and (2) the fraction of the shared TF bindings from the total number of TF binding events, in the promoters of the two paralogs (denoted as “Absolute” and “Relative” similar binding, respectively in **Fig 5C-D** and **Supp Fig 9**). Thus, for each paralog pair, we computed the total number and relative fraction of TF bindings out of all binding events that are shared between the two genes. We plotted the distributions of both the absolute number and relative fraction of shared TFs, for each class of paralogs, partitioned based on their inferred time of duplication, and compared these distributions using a one sided Mann-Whitney test, followed by FDR-correction.

### Promoter sequence conservation analysis of CGI and CGI-less genes

Sequence conservation in promoter regions across mammals was estimated using a 7-way phyloP conservation score, where seven mammals are included(55). An averaged histogram was built for the region encompassing the 1000bps upstream of the TSS (using coordinates of the human gene). This was done separately for the set of CGI and CGI-less genes. Results are shown in **Supp Fig 8**.

### Statistical analysis

The significance of differences in the distributions of rates of gene gain and loss betwen CGI-less and CGI genes (**Fig 1B-C** and **Supp Fig 3**) was computed using a one-sided Mann-Whitney test. For comparisons of human-non human species fractions of one-to-one orthologs between CGI and CGI-less genes (**Fig 1A** and **Supp Fig 2**), a chi-square test was performed on a per-species basis, followed by an FDR-correction.

The distributions of the three classes of paralogs (CGI-CGI, Mixed and CGI-less) across different evolutionary times were analyzed using a permutation test that tested the likelihood of producing a more extreme distribution of CGI-CGI paralogs among older paralogs (**Fig 2** and **Supp Fig 4**). This was done by summing up the CGI-CGI paralog numbers from the most ancient paralogs up to each discrete branch point (or alternativity, to a dS bin) and comparing the observed partition to a random shuffle of the same number of CGI-CGI, Mixed and CGI-less – CGI-less paralogs. A permutation was considered more extreme if it yielded a higher number of CGI-CGI pairs for at least a single branch point (or a dS bin). In each case, 100,000 random shuffling events were considered to obtain an empirical P-value. In humans, in the case of the dS-bins based analysis, for CGI promoters defined using the ‘relaxed’ approach, we only considered the first 23 dS-bins and excluded the 24th. This was because of a relatively large group of CGI-CGI paralogs that had problematic dating of duplication time (based on the two inference methods) that inflated the numbers of CGI paralogs in that bin and were skewing the test.

Transcriptional divergence and similarity of TF-binding of CGI-CGI pairs were compared with CGI-less – CGI-less pairs using one-sided Mann-Whitney tests followed by FDR-corrections (**Figs 3**,**5** and **Supp Figs 5**,**9**).

For TF analysis of human and mouse CGI and CGI-less genes using the Cistrome data, where the TF binding distribution is shown along the TSS coordinates of CGI and CGI-less genes (**Fig 5A-B** and **Supp Fig 7**), it was assumed that the profile of TF binding along the promoter approaches a normal distribution. The comparisons between average CGI and average CGI-less TF-binding profiles were evaluated using a t-test. The test was also performed to compare normalized CGI-less gene profiles between human and mouse, as well as normalized CGI-gene profiles between human and mouse. Statistical tests were done using either the scipy package version 1.5.3(56) or using R (version 4.0.5).

## Acknowledgements

We thank Xi Chen, Peng He, Sivan Friedman-Nakar, Stefan Kaltenbach, Natalia Kunowska, Tomás Pires de Carvalho Gomes, Adi Stern and Michelle Ward for helpful discussions during the analysis and for comments on the manuscript. This research was supported by the Israel Science Foundation (ISF, grant No. 435/20) and by the United States -Israel Binational Science Foundation (BSF, grant No 2019037).

