## Supporting Information for "Promoter architecture links gene duplication with transcriptional divergence"

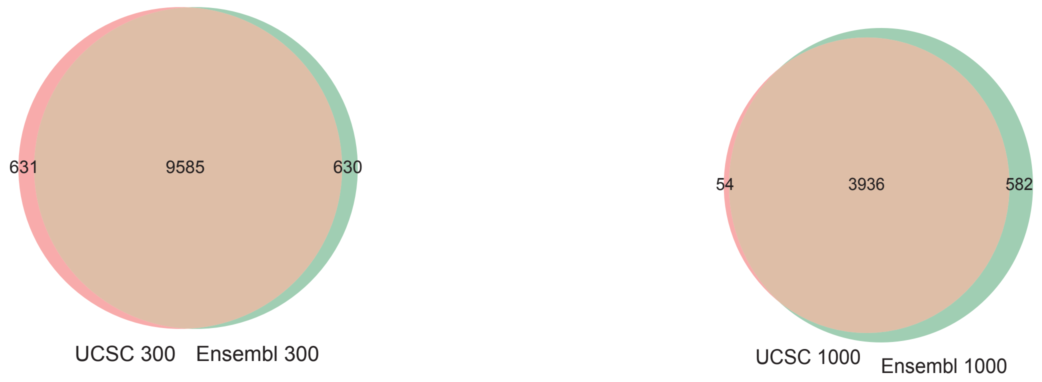


**Supporting Figure 1: Agreement of CGI annotations between UCSC and ENSEMBL.**
Different sources of annotations of CGIs yield similar CGI gene sets: Annotated CGI regions from ENSEMBL and from UCSC were overlapped with promoter regions of genes (regions of 300bp upstream and 100bp downstream of the TSS – **left figure**, or regions of 1,000bp upstream and 100bp downstream of the TSS – **right figure**). Genes were defined as CGI genes if their promoter region overlapped with an annotated CGI for at least 50% of its length. The results, using both promoter lengths, show that the CGI annotations from UCSC and ENSEMBL give similar CGI gene sets (88.4% and 86.1% of the genes are shared in the 300 and 1,000bp versions, respectively).

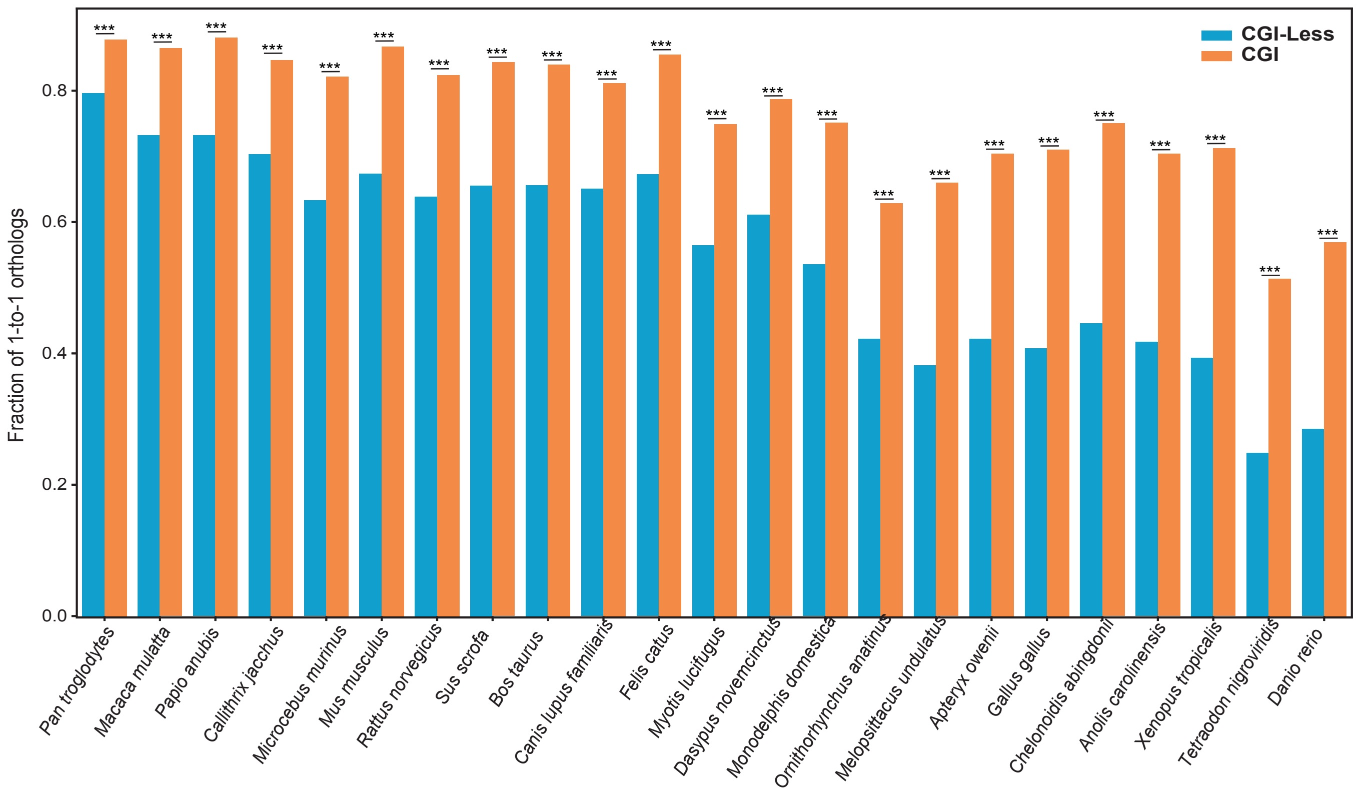


**Supporting Figure 2: CGI and CGI-less genes duplication across species.**
Fraction of 1-to-1 orthologs of human CGI and CGI-less genes (as defined using the inclusive approach described in the main text) with a selected number of species. In each pair of species, the fractions of CGI and CGI-less were compared (the size of group of CGI genes having orthologs, out of the group of all CGI genes, versus the size of group of CGI-less genes having orthologs, out of all CGI-less genes) using a chi-square test. P-values were corrected by FDR (*** - $P<0.001$). In this analysis, CGI genes are identified by the ‘relaxed’ method of overlap of at least 1bp of an annotated CGI region with the 1,000bp up and downstream of the TSS (see analogous analysis with the conservative approach in Fig 1A).

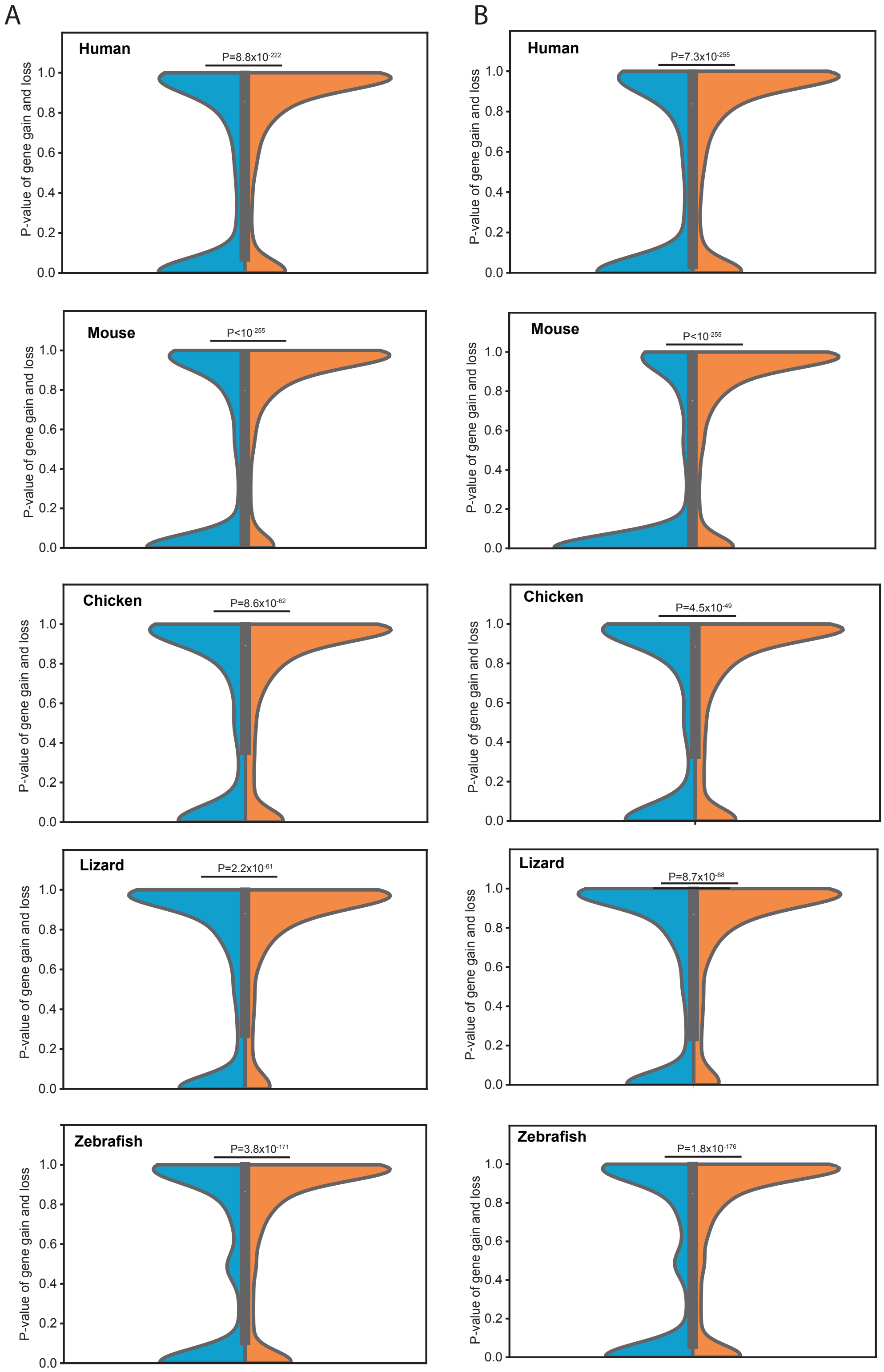


**Supporting Figure 3: CGI and CGI-less genes rates of gain and loss in five vertebrates.**
(**A**) Distribution of P-values of rates of gain and loss in CGI and CGI-less genes in human, mouse, chicken, anole lizard and zebrafish. Comparison between the distributions was preformed using a one-sided Mann-Whitney test. CGI determination was done using the conservative approach. (**B**) Similar to (A), with CGI determination performed using the inclusive approach. In all species and in both methods of CGI determination, CGI-less genes have a distribution enriched with low P-values, suggesting a higher rate of gene gain and loss in this group in comparison with CGI genes.

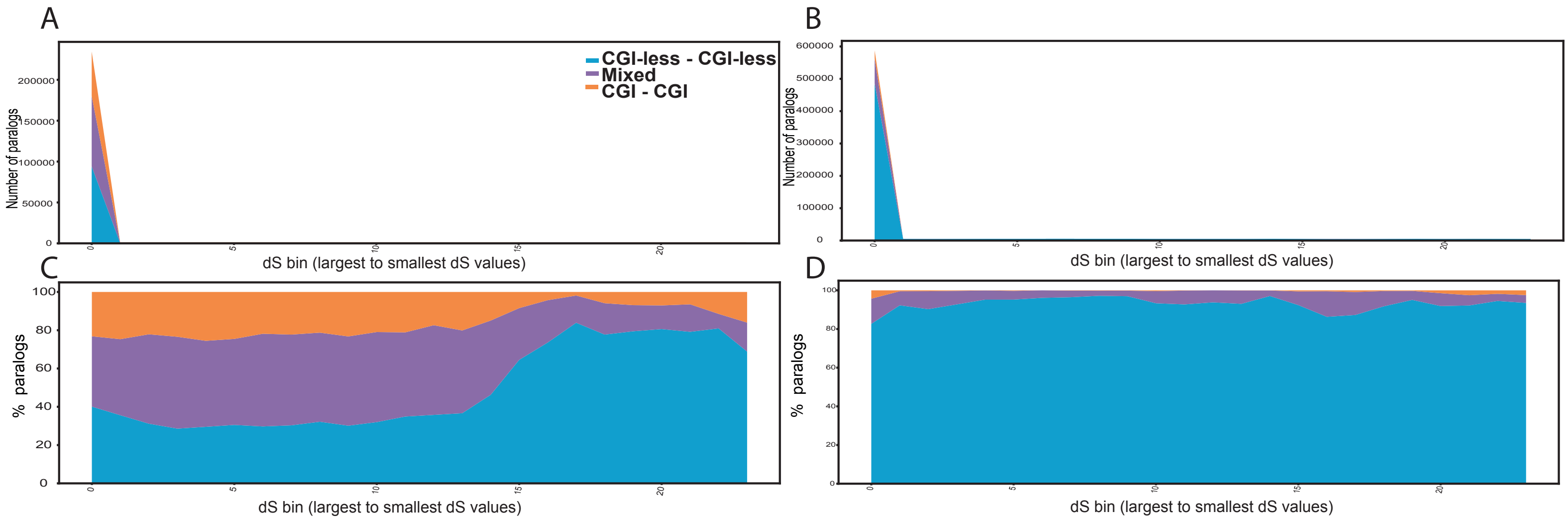


**Supporting Figure 4: Evolutionary timeline of duplication events in CGI and CGI-less genes.**
(**A**) A timeline showing the total numbers of human paralogs that are inferred to have duplicated in different evolutionary time points, split based on the rate of synonymous substitutions - dS (where each paralog pair is binned into one of 24 bins, the left-most having the largest dS values and accordingly enriched with the most ancient paralogs). Paralog pairs are further split based on their promoter classification: CGI-CGI paralogs (orange), CGI-less – CGI-less paralogs (blue) and Mixed (purple). CGI-CGI and Mixed pairs are skewed towards ancient times of duplication ($P\text{-value}<10^{-5}$, permutation test). (**B**) Similar to (A), with mouse paralogs. CGI-CGI and Mixed pairs are skewed towards ancient times of duplication ($P\text{-value}=10^{-5}$, permutation test). (**C**) A timeline showing the relative fractions of human paralogs binned by their dS and split into CGI-CGI, CGI-less – CGI-less, and Mixed paralogs. (**D**) Similar to (B), with mouse paralogs.

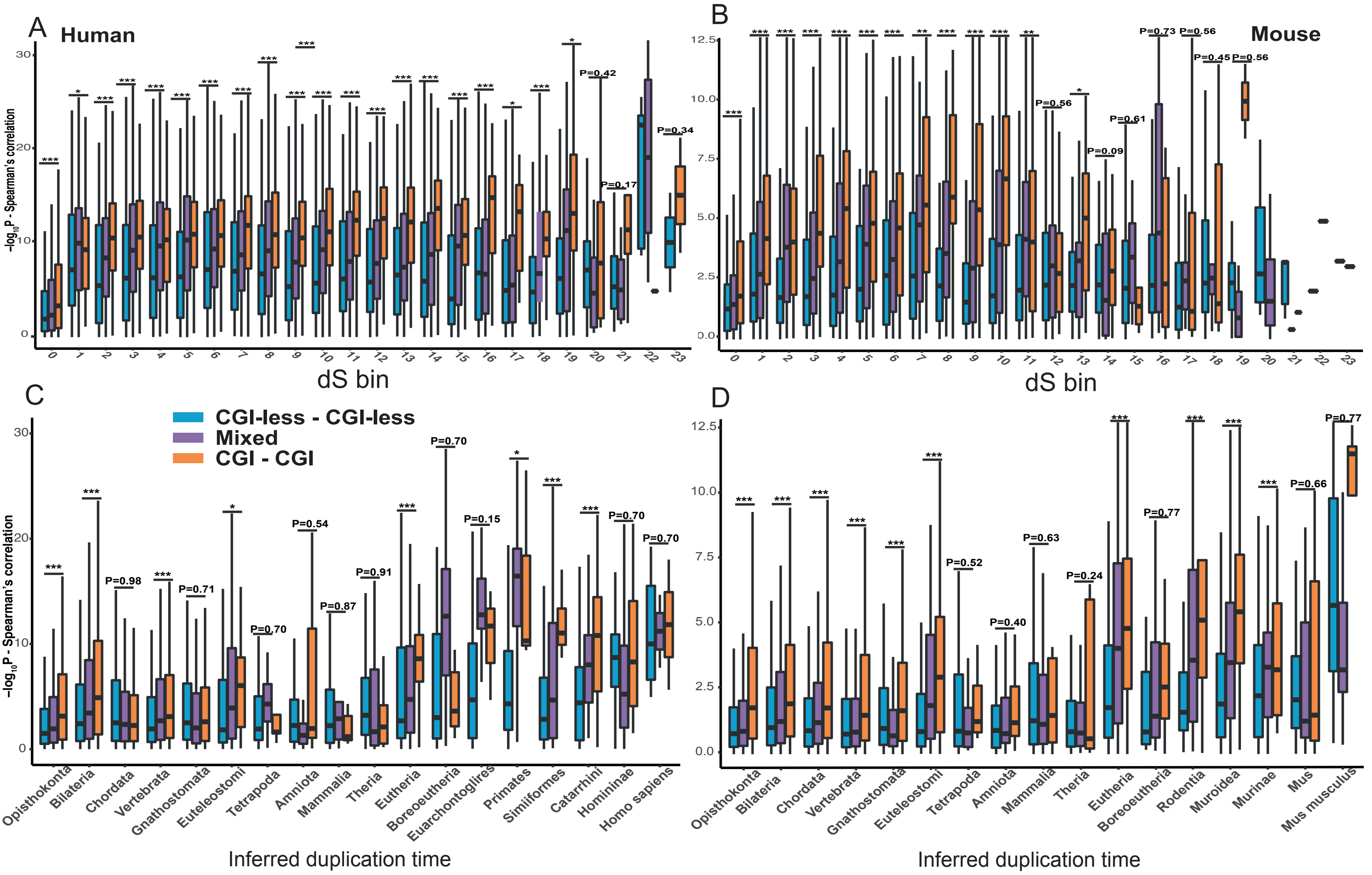


**Supporting Figure 5: Transcriptional divergence in CGI and CGI-less paralogs.** Similar to Fig 3, with Mixed group of paralogs shown. A timeline showing the transcriptional divergence of human (**A,C**) and mouse (**B,D**). Each paralog is classified based on its inferred duplication time. Paralogs are further partitioned to CGI-CGI and CGI-less – CGI-less pairs. Transcriptional divergence is computed based on Spearman's correlation between the expression levels of the two paralogs across tissues and represented using the negative logarithm of the P-value of this correlation – a higher value represents a stronger agreement in expression and a higher correlation in transcription between the two paralogs. In **A-B,** paralogs are partitioned based on bins with similar values of synonymous substitutions (dS), with the lowest bin representing the most diverging paralog set (highest dS values). In **C-D,** the X-axis represnts the inferred duplication time based on tree topology. In each time point, the transcriptional diveregnce of CGI-CGI paralogs is compared with the CGI-less – CGI-less paralogs using a one-sided Mann-Whitney test. FDR-corrected P-values are shown. Groups of duplicates of the same inferred duplication times or dS bins, with a small number of genes were removed in A-D. For clarity, only CGI-CGI and CGI-less – CGI-less paralogs are shown. (***P < 0.001, **P < 0.01, *P < 0.05)

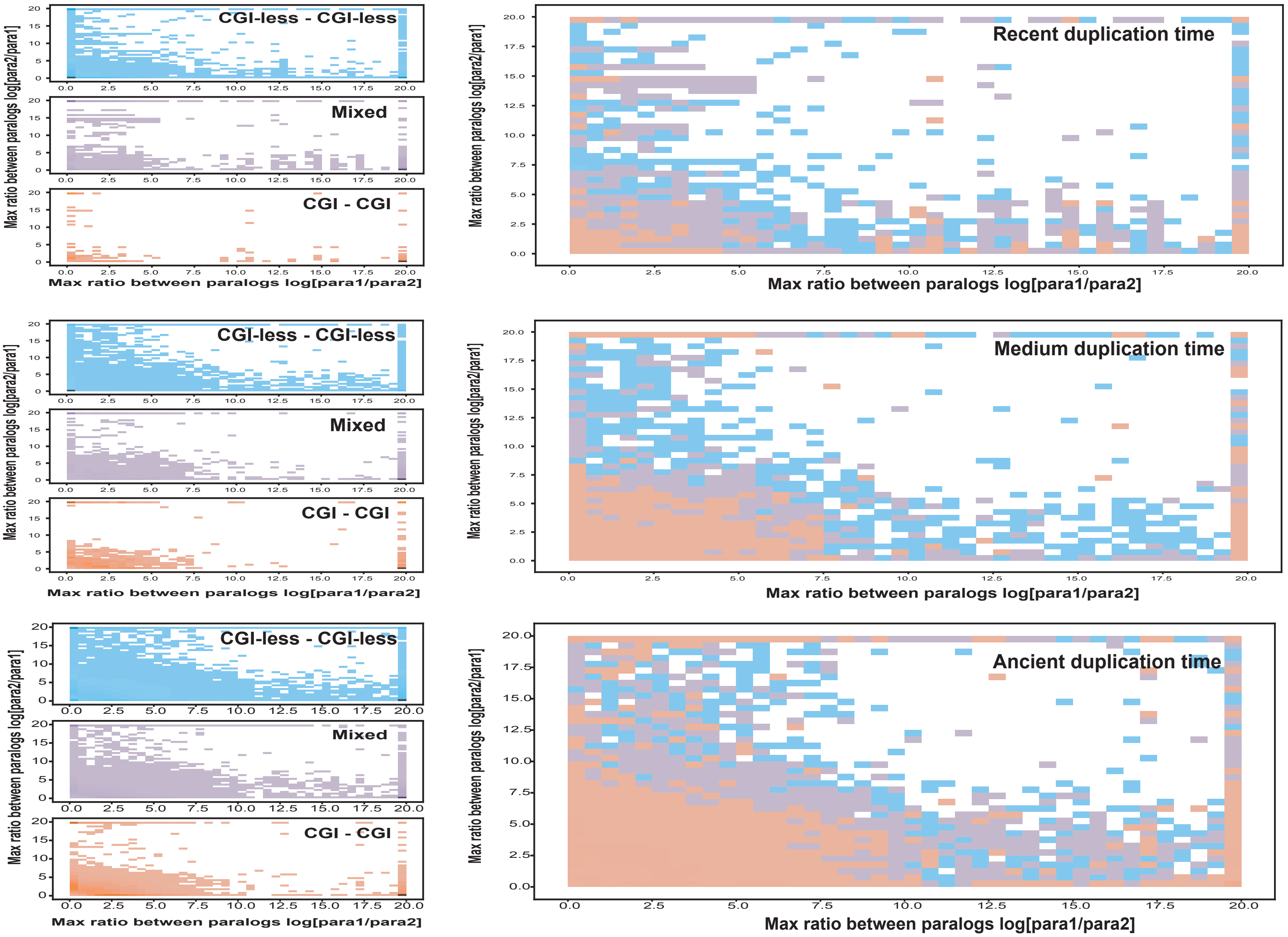


**Supporting Figure 6: Transcriptional sub/neo-functionalization in mouse CGI and CGI-less paralogs.**
As in Figure 4A, with mouse paralogs: Heatmaps showing the log ratio of expression of mouse paralogs, where the X-axis represents the highest ratio between paralog1 and paralog2, and the Y-axis represents the highest ratio between paralog2 and paralog1 (chosen from the set of ratios of expression across tissues). Paralogs are split into recent, medium and ancient duplication times and to CGI-CGI, CGI-less – CGI-less and Mixed paralog classes. Left panels show the three classes separately, right panels show the aggregation of all classes together. Paralog pairs that appear in the top right corner tend to have more distinctive patterns of expression, where each of the pair's paralogs is strongly expressed in a different tissue.

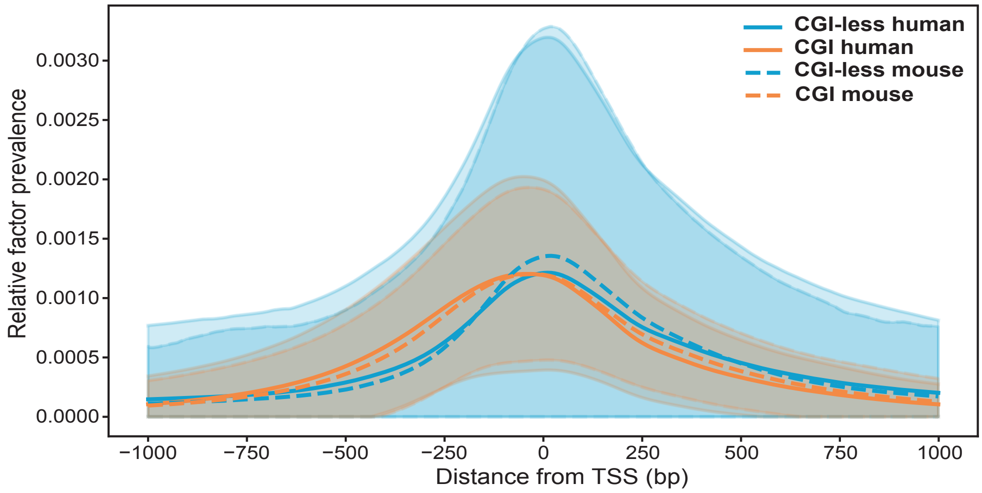


**Supporting Figure 7: normalized TSS-relative histogram of TFs along CGI and CG-less gene promoters.**
For every base pair around the TSS (from 1,000bp upstream to 1,000bp downstream), the TF occupancy is computed by as the total number of recorded TF-ChIP-seq events that intersected it, divided by the total number of genes used in the analysis followed by further normalization to obtain an area of 1 for all curves. CGI and CGI-less gene sets are shown separately (in orange and blue, respectively). The shaded orange and blue regions represent one standard deviation from the average of all CGI and CGI-less genes, respectively. In both human and mouse CGI genes have a wider occupancy around the TSS in comparison with CGI-less genes, which have sharper occupancy (in agreement with previous results(Carninci et al. 2006)). Furthermore, in both human and mouse CGI genes have a slower decay of TF occupancy upstream of the TSS in comparison with the CGI-less gene set. The differences between CGI and CGI-less genes are significant (t-statistics=-48.48, P-value<$1x10^{-130}$X in human, and t-statistics=-39.55, P-value<$1x10^{-130}$ in mouse, t-test). The differences between the CGI and CGI-less genes are more significant in comparison with the differences between human and mouse CGI genes (t-statistics=21.12, P-value<$1x10^{-130}$) or with the differences human and mouse CGI-less genes (t-statistics=8.12, P-value<$1x10^{-130}$), as can be observed by the t-statistics.

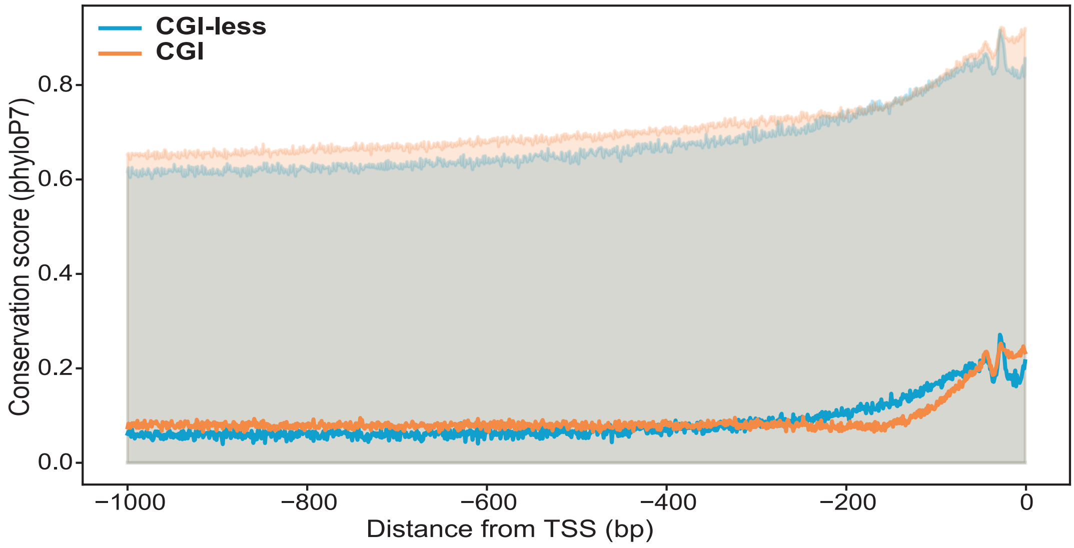


**Supporting Figure 8: Promoter sequence conservation across orthologs of CGI and CGI-less genes.**
Sequence conservation in promoter regions across mammals was estimated using 7-way phyloP conservation scores(Pollard et al. 2010). An averaged histogram was built for the 1,000bp upstream of the TSS for human CGI and CGI-less genes (in orange and blue, respectively). Higher values of phyloP score denotes higher conservation. The orange and blue shaded regions represent one standard deviation from the mean of all human CGI and CGI-less ,respectively. While the averaged data is noisy, it points to several trends suggested before: (1) In innate immune genes, promoter sequence conservation is higher in some regions upstream of the TSS(Schroder et al. 2012, Hagai et al. 2018); (2) In mammals, “Broad promoters”, which are often associated with CGI, differ in their sequence conservation from “Sharp promoters”, which are depleted of CGI regions. The latter group of “Sharp promoters” has a slightly higher conservation in the proximate region of the TSS, in comparison with “Broad promoters”(Carninci et al. 2006). Further to this, the results in Supp Fig 7 may suggest that the higher conservation of CGI-less promoter regions in the immediate proximity to the TSS may be related to the differences in the TF occupancy between CGI and CGI-less genes: While CGI genes have a broader distribution of TF binding which may be more flexible, CGI genes have an effectively narrower region of TF binding nearby the TSS, which may impose stronger constraints on the sequence of this region.

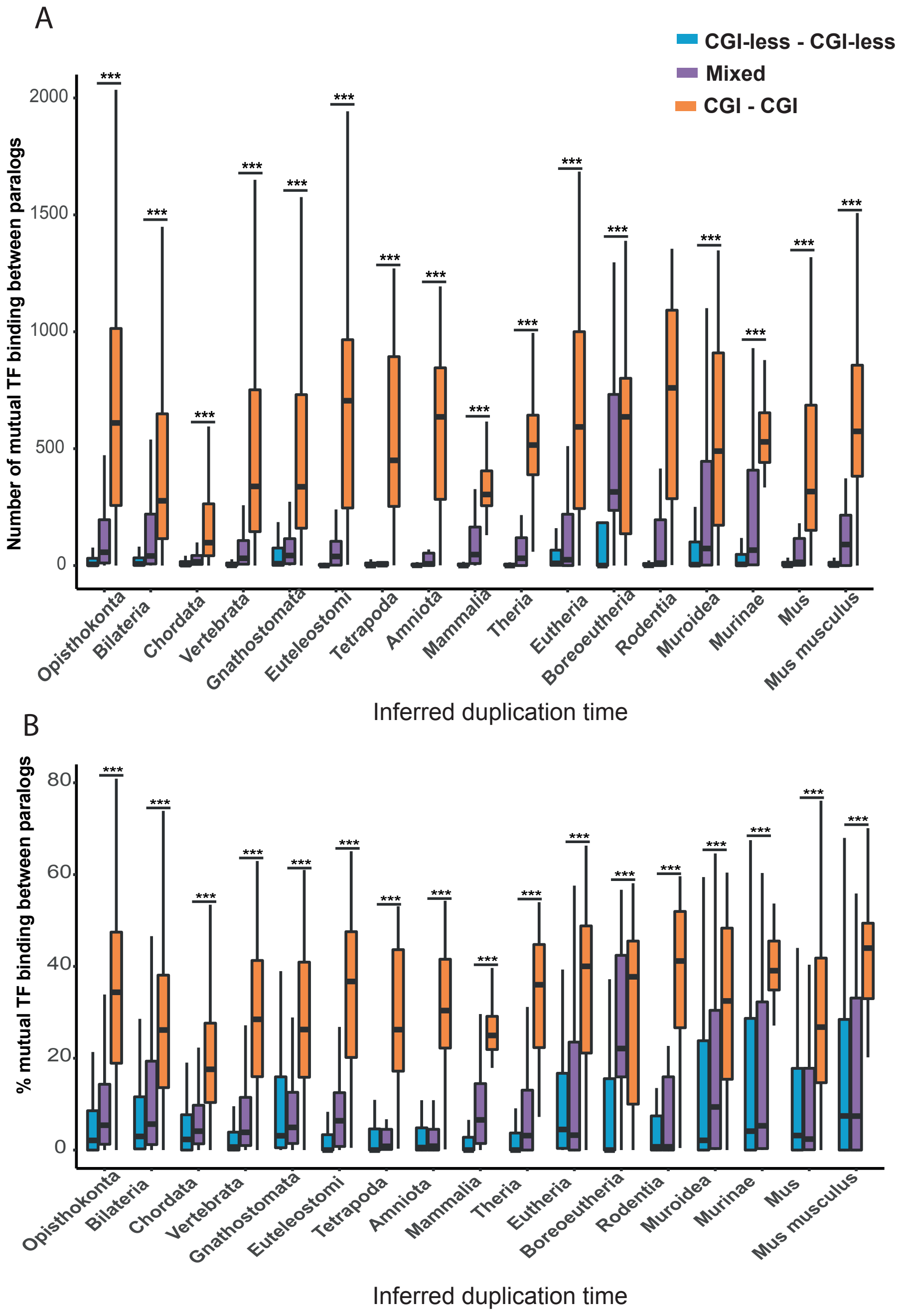


**Supporting Figure 9: Shared TF-binding in promoters of mouse CGI and CGI-less genes and paralogs.**
Similar to Fig 5C-D, in mouse: (**A**) Total number of mutual TF binding to the promoters of CGI-CGI, CGI-less – CGI-less or Mixed paralogs across different inferred duplication times. In each time point, the distributions of the values between CGI-CGI and CGI-less – CGI-less paralogs are compared using a one-sided Mann-Whitney test and corrected by FDR. (**B**) Relative fraction of mutual TF binding to the promoters of CGI-CGI, CGI-less – CGI-less or Mixed paralogs across different inferred duplication times. In each time point, the distributions of the values between CGI-CGI and CGI-less – CGI-less paralogs are compared using a one-sided Mann-Whitney test and corrected by FDR. In all time points, the total number (C) and the relative fraction (D) of the shared TFs between paralogs are significantly larger in CGI-CGI in comparison with CGI-less – CGI-less paralogs. Analogous analyses in mouse give similar results (Supp Fig 9). (***P < 0.001, **P < 0.01, *P < 0.05)
